# Aberrant centrosome biogenesis disrupts nephron progenitor cell renewal and fate resulting in fibrocystic kidney disease

**DOI:** 10.1101/2023.04.04.535568

**Authors:** Tao Cheng, Chidera Agwu, Kyuhwan Shim, Baolin Wang, Sanjay Jain, Moe R. Mahjoub

## Abstract

Mutations that disrupt centrosome structure or function cause congenital kidney developmental defects and fibrocystic pathologies. Yet, it remains unclear how mutations in proteins essential for centrosome biogenesis impact embryonic kidney development. Here, we examined the consequences of conditional deletion of a ciliopathy gene, *Cep120*, in the two nephron progenitor niches of the embryonic kidney. *Cep120* loss led to reduced abundance of both metanephric mesenchyme and ureteric bud progenitor populations. This was due to a combination of delayed mitosis, increased apoptosis, and premature differentiation of progenitor cells. These defects resulted in dysplastic kidneys at birth, which rapidly formed cysts, displayed increased interstitial fibrosis, and decline in filtration function. RNA sequencing of embryonic and postnatal kidneys from Cep120-null mice identified changes in pathways essential for branching morphogenesis, cystogenesis and fibrosis. Our study defines the cellular and developmental defects caused by centrosome dysfunction during kidney development, and identifies new therapeutic targets for renal centrosomopathies.

**Graphical Abstract:** 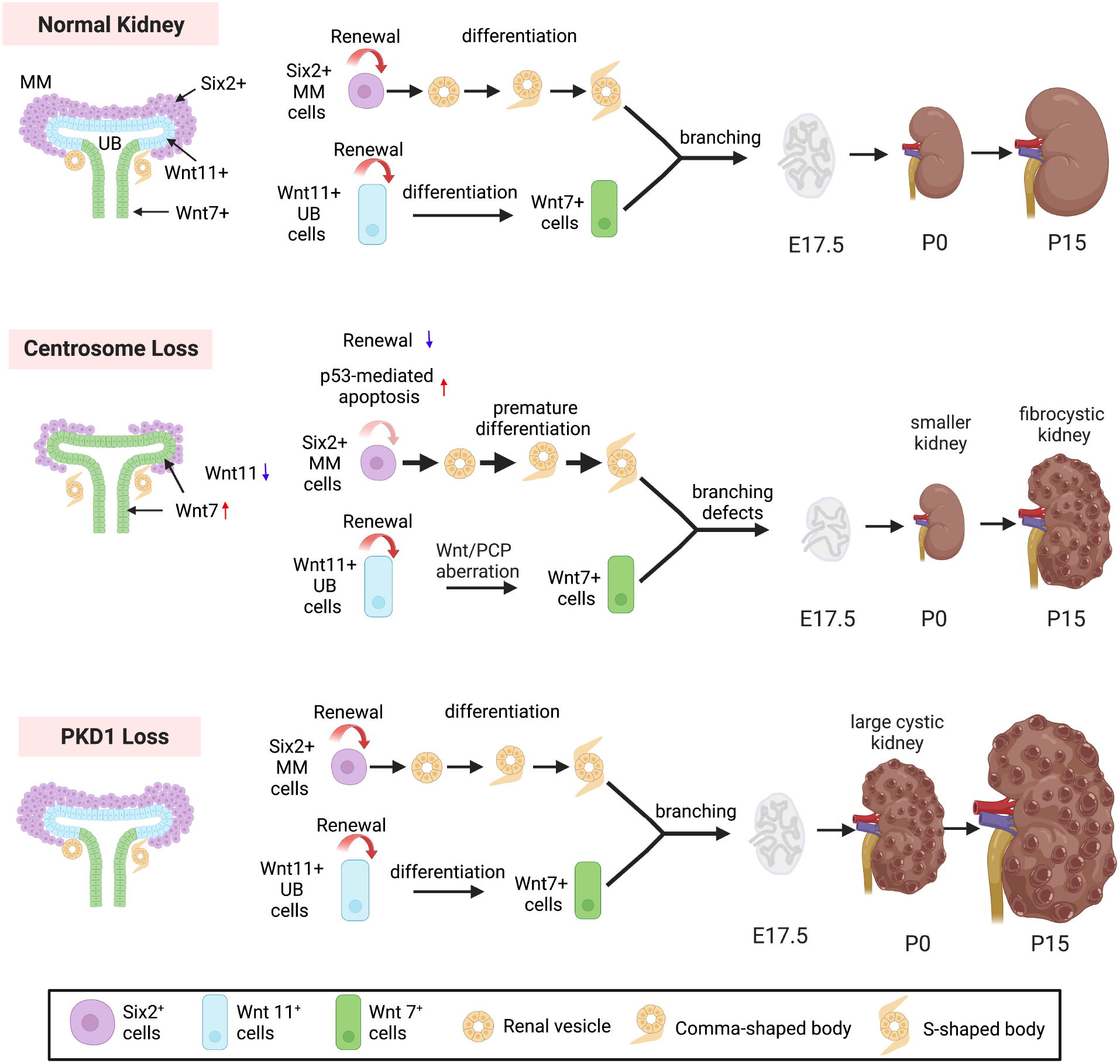

**Highlights:** Defective centrosome biogenesis in nephron progenitors causes:

- Reduced abundance of metanephric mesenchyme and premature differentiation into tubular structures
- Abnormal branching morphogenesis leading to reduced nephron endowment and smaller kidneys
- Changes in cell-autonomous and paracrine signaling that drive cystogenesis and fibrosis
- Unique cellular and developmental defects when compared to Pkd1 knockout models

## Introduction

Mammalian kidney development is a highly coordinated process that requires reciprocal signaling between the metanephric mesenchyme (MM) and ureteric bud (UB) progenitor cells, which is critical for their self-renewal, proliferation, and subsequent differentiation ^1-4^. The UB progenitors undergo extensive growth and several generations of branching morphogenesis to establish the collecting duct system of mature nephrons. At each branching event, the MM cells similarly proliferate at the UB tips in the nephrogenic zone. The progenitors then begin to differentiate, form a tight cluster of cells called the pretubular aggregate that is located beneath the UB branch tip, then epithelializes to form a renal vesicle (RV). The RVs subsequently undergo a series of morphological changes and patterning to form the curved pretubular structures termed the comma- and S-shaped body. These ultimately give rise to the proximal segments of mature nephrons including glomeruli, proximal, and distal tubules ^2,4^. Mutations that disrupt the reciprocal signaling between these two progenitor niches negatively impact their growth and differentiation, leading to defective branching morphogenesis, abnormal nephron development and endowment, congenital renal dysplasia, and early-onset fibrocystic kidney diseases ^1-4^.

A significant number of genes that are mutated in patients with cystic and fibrotic renal pathologies encode cilia- and centrosome-associated proteins ^5-7^. The centrosome is the major microtubule-organizing center in mammalian cells, with important roles in regulating the microtubule cytoskeleton in interphase and mitosis ^8,9^. Cilia are hair-like organelles that are templated by the centrosome and protrude from the apical surface of most cells, including the majority of cell types in the kidney ^10,11^. Together, the centrosome-cilium complex act as a signaling hub to regulate cell-cell communication, cell proliferation, differentiation, fate determination and ultimately tissue formation. Mutations in several centrosome-ciliary genes cause kidney diseases including Autosomal Dominant Polycystic Kidney Disease (ADPKD), Autosomal Recessive Polycystic Kidney Disease (ARPKD), and Nephronophthisis (NPH) ^12-14^. In addition, mutation in these organelles cause multi-organ disease syndromes called ciliopathies, which include Joubert syndrome (JS) and Jeune Asphyxiating Thoracic Dystrophy (JATD) ^15^. Of note, ciliopathy patients develop kidney phenotypes that can vary dramatically. For example, ADPKD is characterized by slow formation of cysts that leads to progressive enlargement of the kidneys, damaging of the renal parenchyma causing fibrosis, and a gradual decline in kidney function leading to end-stage kidney disease (ESKD) in the 5-6^th^ decade of life ^16,17^. In contrast, NPH is a rare earl-onset renal ciliopathy which typically causes ESKD in children and young adults. Unlike ADPKD, renal cysts are not a hallmark of NPH, as they remain small and are usually located at the corticomedullary junction. NPH is characterized by smaller, hyperechogenic kidneys with cortico-medullary cysts and poor cortico-medullary differentiation, tubular atrophy, disintegration, and irregular thickening of the tubular basement membrane, leading to fibrotic kidneys with shrunken appearance ^18^. Similarly, JS and JATD patients commonly display abnormal congenital kidney developmental phenotypes such as dysplastic, fibrocystic kidneys and reach ESKD within childhood or adolescence ^19,20^. So far, ciliopathies have been linked to mutations in ∼ 200 genes ^6,7^, a significant number of which encode centrosomal proteins, and many of these mutations result in congenital kidney developmental defects and early-onset fibrocystic disease phenotypes. This suggests a critical role for centrosomes in the development of embryonic kidneys.

The centrosome is comprised of a pair of centrioles surrounded by a mesh of proteins called the pericentriolar material. The older of the two centrioles templates the assembly of the cilium. Centrosomes are duplicated once every cell cycle and segregated to each daughter cell following mitosis, and this process is tightly controlled such that each cell inherits a single centrosome and forms a single cilium ^21,22^. Mutations in genes that disrupt the centrosome duplication cycle can result in daughter cells that contain aberrant centrosome numbers, which in turn lead to pathological phenotypes ^23^. Specifically, mutations in centriole duplication factors can block centrosome biogenesis and result in cells lacking centrosomes after several rounds of cell division, a phenomenon termed **C**entrosome **L**oss (CL) ^24,25^. Several cellular and molecular changes occur following CL. Since centrosomes facilitate the assembly of the mitotic spindle, cell cycle progression and mitosis are often impaired in cells lacking centrosomes ^24,25^. CL in wildtype cells leads to prolonged mitosis, p53-dependent cell cycle arrest, activation of the p53BP1-USP28-TP53-dependent mitotic surveillance mechanism, and caspase-mediated apoptosis ^24,25^. Inducing CL globally in mice causes prometaphase delay and p53-dependent apoptosis in most cells, which prevents embryo development upon midgestation, and causes lethality by embryonic day nine ^26^. Conditional induction of CL in the developing mouse brain leads to neural progenitor cell death and premature differentiation into neurons, ultimately leading to microcephaly phenotypes ^27^. Inducing CL in the developing lung endoderm similarly causes p53-mediated apoptosis, but only in proximal airway cells with low extracellular signal-regulated kinase (ERK) activity ^28^. Meanwhile, centrosomes of the intestinal endoderm appear to be dispensable for progenitor cell growth and differentiation during development. Thus, the consequences of CL on progenitor cell growth and fate differs depending on the specific type of the cell, tissue, and developmental context.

Centrosomal protein 120 (Cep120) is a daughter centriole-enriched protein that plays essential roles in centriole duplication, elongation, and maturation ^29-31^. Previous studies have shown that depletion of Cep120 in proliferating cells result in defective centriole biogenesis and loss of centrosomes after two cell divisions ^29-31^. In addition, Cep120 depletion in non-dividing quiescent cells disrupts centrosome homeostasis, causing excessive pericentriolar material accumulation, defective microtubule nucleation and dynein-dependent cargo trafficking, ultimately leading to aberrant ciliary assembly and signaling ^32^. Genetic ablation of Cep120 globally in mice results in early embryonic lethality ^30^, while siRNA-mediated depletion or conditional deletion of the gene in cerebellar granule neuron progenitors (CGNPs) causes hydrocephalus and cerebellar hypoplasia ^30,33^. These phenotypes are due to failed centrosome duplication, aberrant microtubule nucleation, abnormal maturation of CGNPs, and defective ciliogenesis in ependymal cells ^30,33^. Importantly, mutations in Cep120 were recently shown to cause JATD and JS, with patients showing complex ciliopathy phenotypes including severe congenital kidney developmental defects and early-onset fibrocystic pathologies ^34-36^. Yet, it remains unknown how mutations in Cep120, and thus defects in centrosome biogenesis, impact kidney development and cause the fibrocystic kidney disease phenotypes.

In this study, we investigated the consequences of CL on renal progenitor cell growth and differentiation, nephron formation, kidney development and filtration function. We show that conditional knockout of *Cep120* in the MM or UB progenitor cells during embryonic kidney development blocked centrosome biogenesis and led to reduced progenitor cell abundance. Centrosome loss caused premature differentiation of both progenitor niches, resulting in early formation of pre-tubular structures, decreased branching morphogenesis, and small dysplastic kidneys at birth. Transcriptional profiling of embryonic kidneys from Cep120-KO mice identified changes in several cellular processes regulated by Wnt signaling. Postnatally, the centrosome-less kidneys rapidly developed fibrosis and formed small cysts, which together resulted in a decline in renal function. Finally, transcriptional profiling of postnatal fibrocystic kidneys from Cep120-KO mice identified unique pro-fibrogenic and cystogenic signaling signatures that are involved in cell proliferation, migration, and extracellular deposition.

## Results

### Centrosome loss in nephron progenitors results in small dysplastic kidneys at birth

To disrupt centrosome biogenesis and cause centrosome loss in a spatiotemporally controlled manner in nephron progenitor cells, we used a recently developed mouse model harboring a floxed allele of Cep120 (^30^; Figure 1A). To induce CL in the metanephric mesenchyme (MM), *Cep120^F/F^* mice were crossed with the Six2-Cre strain ^37^ that expresses Cre-recombinase in the MM lineage (hereafter called Six2-Cep120; Figure 1A). To cause CL in the ureteric bud (UB) progenitors, Cep120^F/F^ mice were mated to a Hoxb7-Cre strain ^37^ (hereafter named Hoxb7-Cep120; Figure 1A). To compare the differences between centrosome loss and a ciliary signaling defect in nephron progenitors during embryonic kidney development, we crossed *PKD1^F/F^* mice ^38^ to the same two Cre-recombinase expressing strains (hereafter named Six2-Pkd1 and Hoxb7-Pkd1, Figure 1A). We examined the extent and specificity of Cep120 KO by staining kidney sections from E13.5 mice with antibodies against Cep120. Compared to wild-type control littermates, Cep120 expression was lost in the Six2-positive mesenchyme but still evident in the UB cells of Six2-Cep120 mice (Figure S1A). Similarly, Cep120 signal was missing in E-cadherin–positive UB tips of Hoxb7-Cep120 KO mice, but present in the MM (Figure S1A). Roughly 90% of MM cells had no Cep120 expression, whereas 85% of UB cells had lost Cep120, respectively (Figures S1B and S1C).

**Figure 1.**
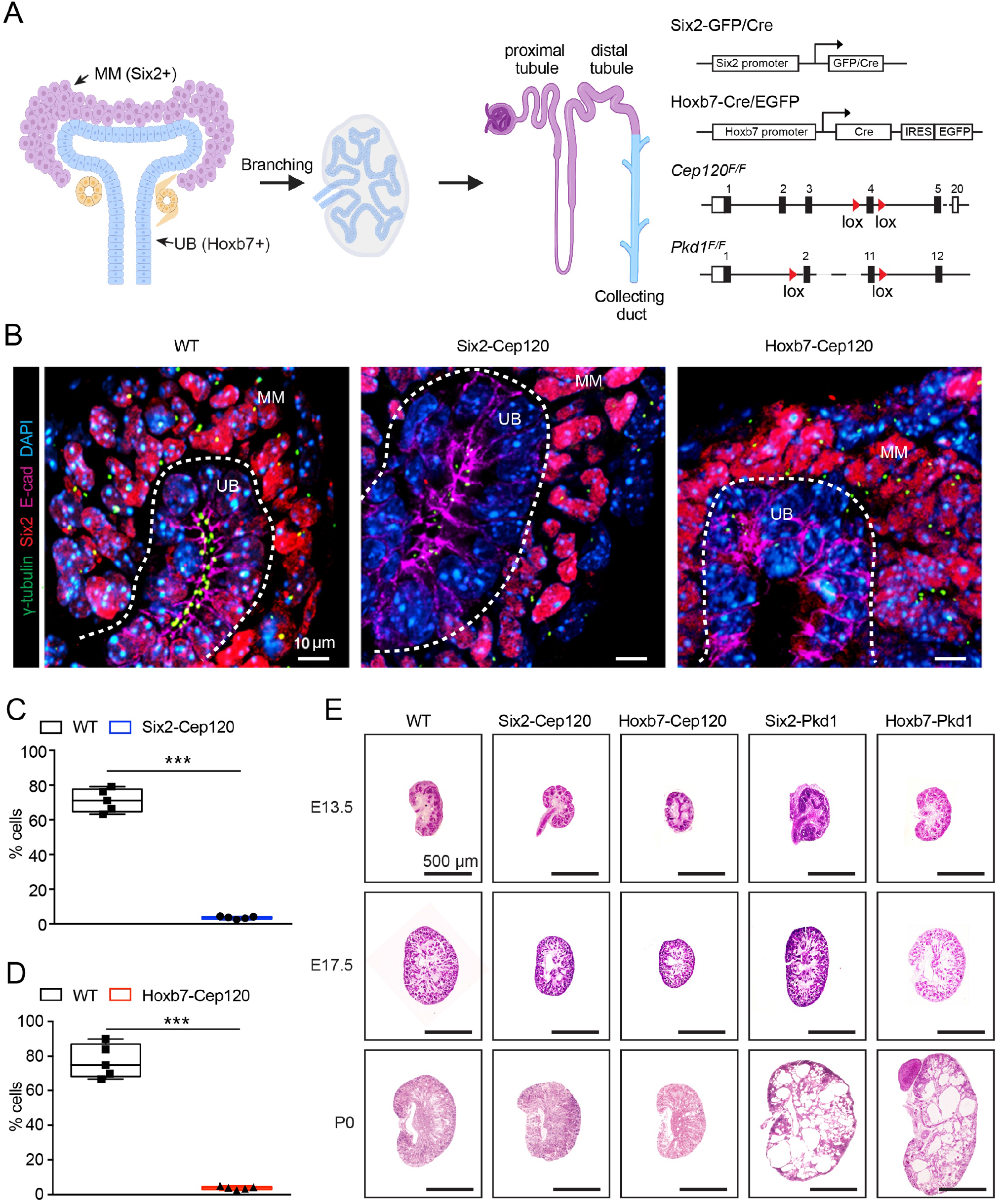
Centrosome loss in nephron progenitors results in small dysplastic kidneys at birth. (A) Schematic representation of nephrogenesis, from renal progenitors to mature nephrons. Diagram (right) highlights genetic cross to conditionally ablate Cep120 or Pkd1 with Six2-Cre (magenta) and Hoxb7-Cre (blue). (B) Immunostaining for centrosomes (γ-tubulin), MM (Six2) and UB (E-cadherin) in kidney sections of E13.5 Six2-Cep120, Hoxb7-Cep120 KO and control mice. (C) Quantification of the percentage of cells lacking centrosomes in the MM population of E13.5 Six2-Cep120 KO and control kidneys. (D) Quantification of the percentage of cells lacking centrosome in UB epithelia of E13.5 Hoxb7-Cep120 KO and control kidneys. (E) H&E staining of kidney sections from various embryonic kidney developmental stages. N = 5 mice per group.

Next, we determined the effect of Cep120 loss on centrosome biogenesis in the two progenitor populations. In Six2-Cep120 kidneys, there was a 71% decrease in centrosome numbers in the MM, whereas centrosome formation appeared normal in neighboring tubular cells (Figures 1B and 1C) Similarly, there was a 77% decrease in centrosome numbers in the UB cells of Hoxb7-Cep120 kidneys, whereas centrosome formation was unaffected in the MM (Figures 1B and 1D). Importantly, there were no changes in Cep120 expression nor centrosome loss in either Hoxb7-Pkd1 or Six2-Pkd1 mice (Figures S1A-S1D). Thus, loss of Cep120 results in cell type-specific block in centrosome duplication during embryonic kidney development in our models.

To determine the consequences of CL on overall kidney morphology, we performed H&E staining of kidney sections isolated at E13.5, E17.5 and P0. Compared with WT controls, kidneys of Six2-Cep120 and Hoxb7-Cep120 mice were smaller in size, which was evident as early as E13.5, and persisted until birth (Figures 1E and 6C). In contrast, both Pkd1 KO mice displayed normal sized kidneys at embryonic stages, which rapidly developed cysts and were significantly enlarged compared to WT at P0 (Figures 1E and 6C). These data indicate that impaired centrosome biogenesis results in unique embryonic kidney developmental defects, which are not evident upon loss of Pkd1 and ciliary signaling function.

### Defective centrosome biogenesis leads to reduced mesenchymal progenitor abundance

To determine how centrosome biogenesis defects during embryonic kidney development cause the small kidney phenotype, we investigated the growth of nephron progenitor cells and their differentiation into pretubular structures. First, we quantified the density of Six2-positive MM at UB tip structures of the nephrogenic zone from kidneys isolated at E13.5. Both Six2-Cep120 and Hoxb7-Cep120 KO kidneys had similar amounts of Six2-positive mesenchymal cells compared with controls at E13.5 (Figure 2C), indicating that centrosome loss had not yet impacted the abundance of Six2-positive cells at this stage. However, the density of Six2-positive mesenchyme was significantly reduced in both KO models beginning at E17.5 (Figures 2A and 2C). Notably, both Six2-Pkd1 and Hoxb7-Pkd1 KO kidneys had similar densities of Six2-positive cells compared to WT at E13.5 and E17.5 (Figures 2A and 2C). These results suggest that centrosome loss and ciliary signaling defects (due to *PKD1* ablation) have distinct effects on mesenchymal progenitor cell growth and renewal.

**Figure 2.**
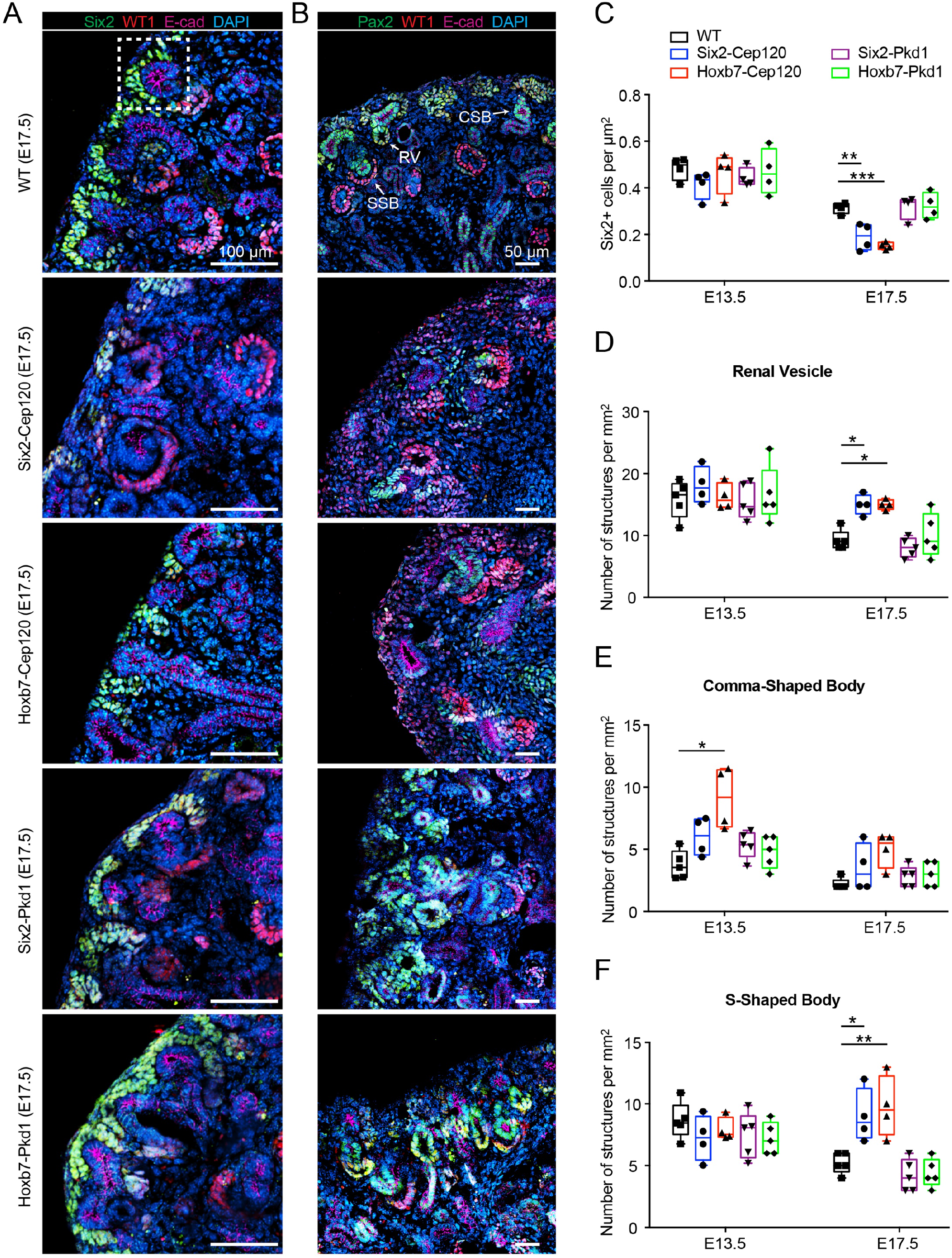
Defective centrosome biogenesis leads to reduced mesenchymal progenitor abundance and premature differentiation. (A) Immunostaining for MM (Six2), pretubular structures (WT1) and UB (E-cadherin) in kidney sections of Six2-Cep120, Hoxb7-Cep120, Six2-Pkd1, Hoxb7-Pkd1 and control mice at E17.5. (B) Immunostaining for pretubular structures (Pax2 and WT1) and UB (E-cadherin) in kidney sections of Six2-Cep120, Hoxb7-Cep120, Six2-Pkd1, Hoxb7-Pkd1, and control mice at E17.5. (C) Quantification of Six2+ cell density in kidney sections of Six2-Cep120, Hoxb7-Cep120, Six2-Pkd1, Hoxb7-Pkd1 KO and control mice at E13.5 and E17.5. N = 4 mice per group. (D-F) Quantification of the density of RV (D), CSB (E) and SSB (F) in kidney sections of Six2-Cep120, Hoxb7-Cep120, Six2-Pkd1, Hoxb7-Pkd1 and control kidneys at E13.5 and E17.5. N = 5 mice (WT, Six2-Pkd1 and Hoxb7-Pkd1) and N = 4 mice (Six2-Cep120 and Hoxb7-Cep120).

There are at least two mechanisms by which the growth of progenitor cells can be disrupted upon defective centrosome biogenesis; (i) CL can result in prolonged mitosis, activation of the mitotic surveillance pathway leading to nuclear translocation of p53 and caspase-mediated apoptosis, ultimately resulting in reduced stem cell proliferation ^24,25,27,39^; (ii) CL has also been shown to disrupt cell fate and cause premature differentiation, depleting the pool of progenitor cells in the brain ^27,28^. To determine which of these processes is impacted in renal mesenchymal cells lacking centrosomes, we first asked whether cells with CL experience prolonged mitosis. This was determined by quantifying the fraction of mitotic cells (phosphorylated Histone H3-positive, pHH3+) among the total pool of Ki67+ cycling progenitors^27^. Both Six2-Cep120 and Hoxb7-Cep120 kidneys showed an increased mitotic index when compared to control kidneys even though the total proliferative population was the same (Figures S2A and S2B), suggesting cells are delayed passaging through mitosis. We next evaluated whether this mitotic delay leads to p53-dependent apoptosis, by staining kidney samples for p53 and cleaved Caspase 3. There was a significant increase in the percentage of cells with nuclear-enriched p53 in both Six2-Cep120 and Hoxb7-Cep120 kidneys (Figures S2C and S2D). The percentage of apoptotic cells (cleaved Caspase 3+) per unit area was also increased in kidneys of both Six2-Cep120 and Hoxb7-Cep120 at E13.5 (Figures S2E-S2F).

Next, we examined the impact of CL on the differentiation capacity of mesenchymal progenitors into nephron tubules. Nephron formation commences when a few Six2-positive MM cells at a new branch tip undergo mesenchymal-to-epithelial transition (MET) and sequential morphological alterations to form a pretubular structure called the renal vesicle (RV), which then develops into comma-shaped body (CSB), S-shaped body (SSB), and ultimately form the proximal segments of the mature nephron ^1^ (Figure 1A). Quantification of RV and SSB density in Six2-Cep120 and Hoxb7-Cep120 kidneys showed no difference compared with control mice at E13.5 (Figures 2B, 2D and 2F). However, there was an increase in the abundance of CSB in Hoxb7-Cep120 kidneys evident as early as E13.5 (Figures 2B and 2E), suggestive of potential acceleration in nephron tubular epithelia specification. Indeed, both KO kidneys contained significantly higher levels of RV and SSB structures compared to WT at E17.5 (Figures 2B, 2D and 2F). These results suggest that Six2-positive mesenchyme lacking centrosomes undergo premature differentiation into nephron tubular structures. In contrast, the densities of RV, CSB and SSB in Six2-Pkd1 and Hoxb7-Pkd1 KO kidneys were unchanged compared to WT kidneys at all embryonic stages (Figures 2B-2F). Collectively, these data indicate that centrosome loss causes both cellular apoptosis and premature differentiation of mesenchymal progenitors, which together result in reduced abundance of this stem cell population.

### Centrosome loss disrupts ureteric bud branching morphogenesis

Next, we determined the effects of CL on the growth of ureteric bud progenitor cells and overall branching morphogenesis. Whole kidneys from Six2-Cep120 and Hoxb7-Cep120 KO mice at E15.5 were isolated and optically cleared using the CUBIC protocol, then stained for the ureteric bud epithelium marker E-cadherin. These samples were transparent and permissive for imaging by light-sheet microscopy ^40^ (Figure 3A). Quantification of branch tips and nodes using the Imaris software Filament Analysis tool showed that Six2-Cep120 and Hoxb7-Cep120 KO kidneys contain significantly lower number of branch tips compared with WT kidneys at the same developmental stage (Figures 3B and 3C). In addition, the number of branching nodes were lower compared to WT (Figure 3D). These data point to potential defects in UB progenitor cell growth and renewal, which are essential for continuous branching morphogenesis ^41^. Finally, there was an increase in nephron tubule diameter in Six2-Cep120 and Hoxb7-Cep120 KO kidneys at E15.5 (Figure 3E), suggesting dilations in these tubules occur early during development. Finally, quantification of glomerular number showed a significant decrease in both Cep120-KO models at P0 (Figure S2G), indicative of reduced nephron endowment. In sum, these data indicate that centrosome loss in the UB niche is deleterious for ureteric bud progenitor growth that is required for branching morphogenesis during early embryonic kidney development.

**Figure 3.**
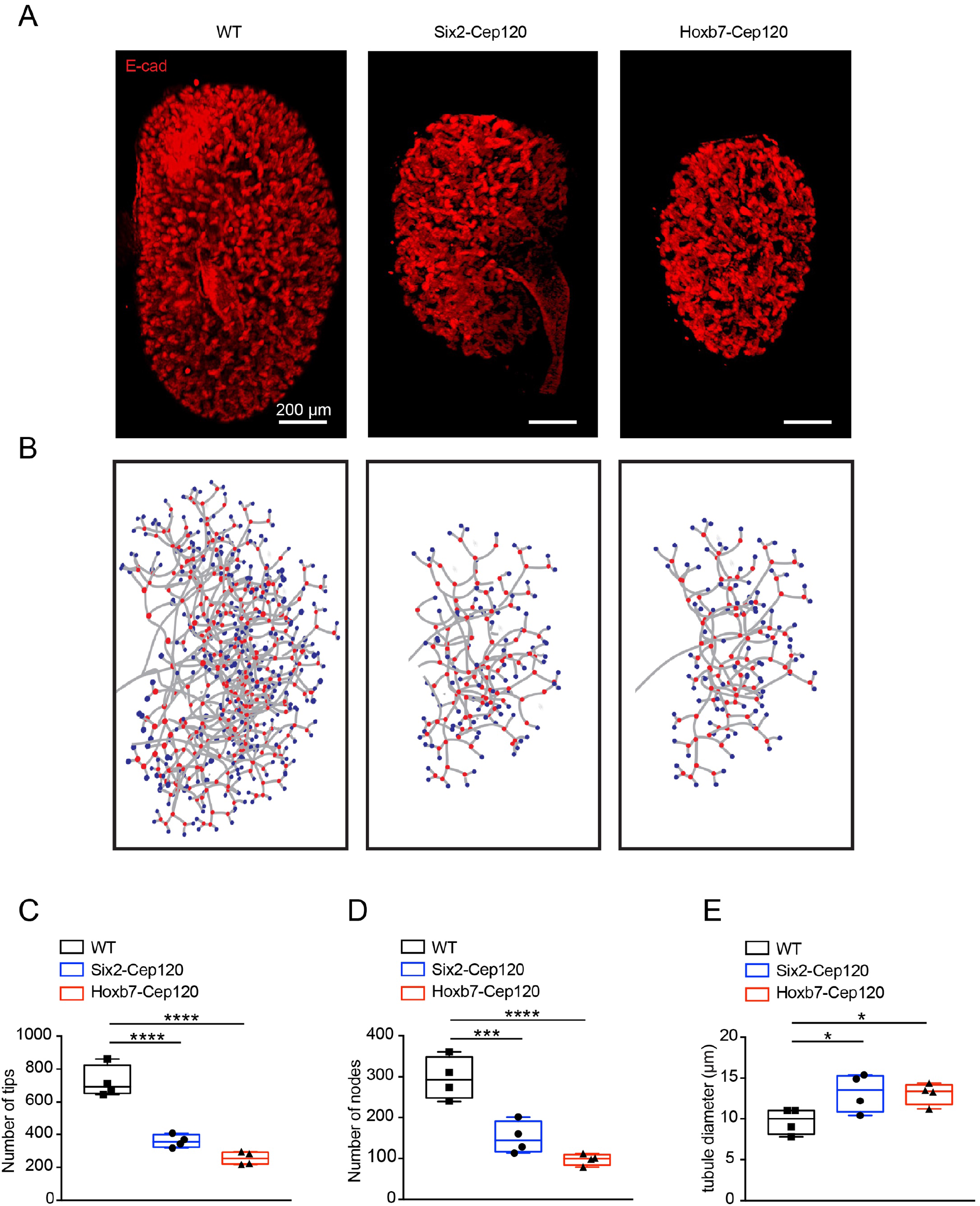
Centrosome loss disrupts ureteric bud branching morphogenesis. (A) Light-sheet microscopy images of whole-mount kidneys from Six2-Cep120, Hoxb7-Cep120 and control mice at E15.5 immunostained for the nephron tubule marker E-cadherin. (B) Imaris filament rendering of ureteric branching in E15.5 kidneys. (C) Quantification of the number of branch tips of Six2-Cep120, Hoxb7-Cep120 KO and control kidneys. (D) Quantification of the number of branch nodes of Six2-Cep120, Hoxb7-Cep120 KO and control kidneys. (E) Measurement of tubule diameter in kidneys of Six2-Cep120, Hoxb7-Cep120 KO and control mice. N = 4 mice per group.

### Transcriptional profiling of centrosome-less cells identifies changes in nephrogenic signaling pathways

To identify pathways that are disrupted upon CL that result in the dysplastic kidney phenotype, we performed transcriptional profiling of centrosome-less embryonic kidneys at the onset of the observed developmental defects. Since the Hoxb7-Cre transgene co-expresses GFP in the ureteric bud lineage ^37^, we isolated UB progenitors from Cep120 KO kidneys and WT control at E13.5 by FACS sorting, followed by bulk RNAseq analysis (Figure 4A, Supplementary Excel 1). Differential expressed gene (DEG) analysis was performed using |log2 fold-change| ≥ 2 and p-value < 0.05. There was a total of 618 DEGs (upregulated: 397; downregulated: 121) in the Cep120-KO UB cells compared to wildtype (Figure 4B). Hallmark pathway analysis identified several key biological and developmental signaling pathways impacted upon centrosome loss (Figure 4C). These include upregulation of mitotic spindle regulators NUMA1, Cdk5rap2, and PCNT, which play critical roles in organizing the mitotic spindle poles and controlling spindle orientation in the absence of centrosomes ^42-46^. Moreover, there was upregulation of components of the TNFα-NF-κB, myogenesis, UV stress response, hypoxia and TGFβ signaling pathways. In conjunction, there was downregulation in oxidative phosphorylation (OXPHOS) and fatty acid metabolism (FAO) (Figure 4C). Together, these alterations suggest that centrosome loss likely causes oxidative stress, activation of pro-inflammatory signaling and mitochondrial defects resulting in tubular injury and inflammation that can drive fibrosis ^47-50^.

**Figure 4.**
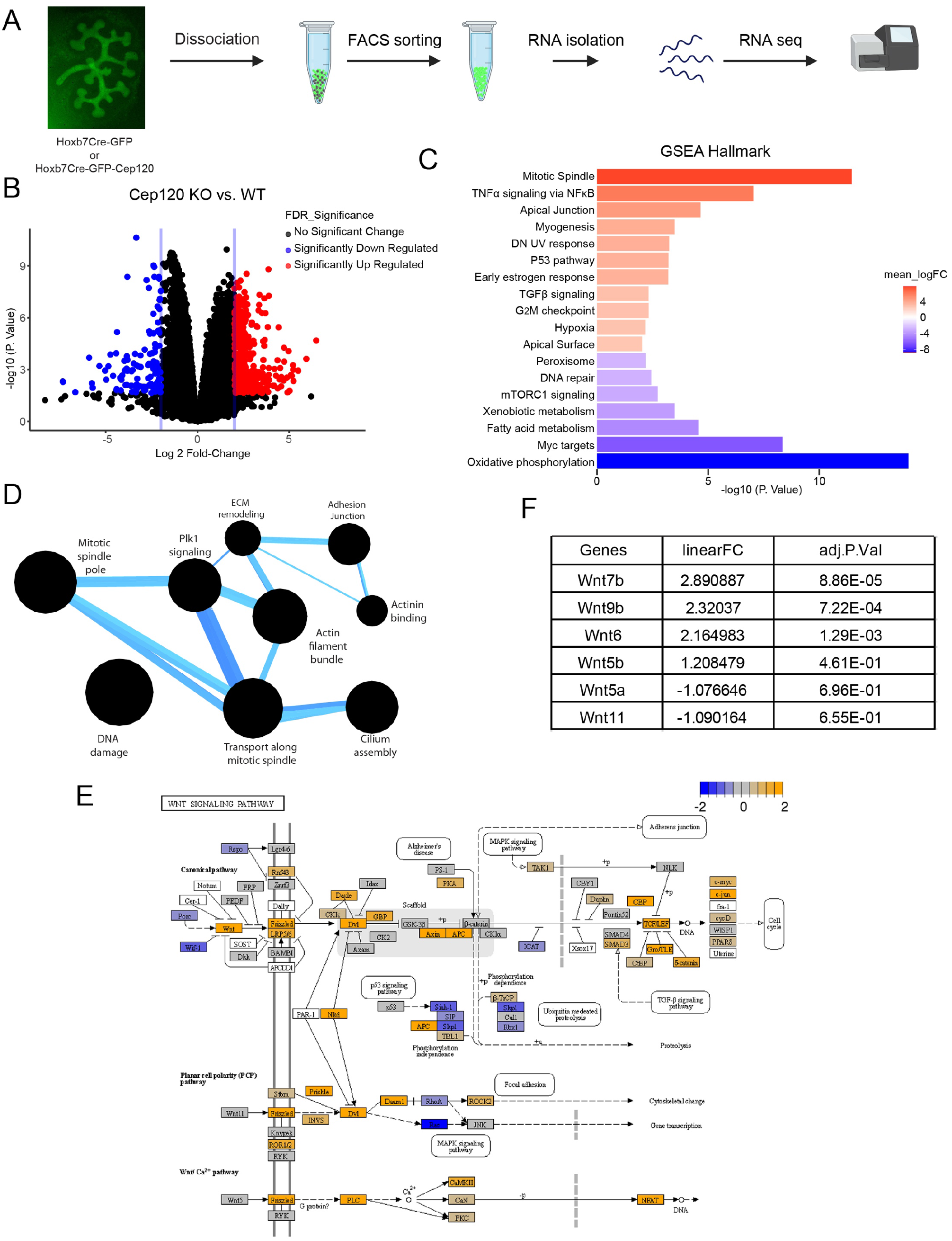
Bulk RNA sequencing analysis of Hoxb7-Cep120 KO and control kidneys. (A) Schematic representation of GFP+ cell sorting and bulk RNA-seq analysis of control and Hoxb7-Cep120 kidneys at E13.5. (B) Volcano plot showing differentially expressed genes in Hoxb7-Cep120 KO versus control kidneys at E13.5. N = 3 mice per group. (C) Hallmark pathway analysis of Hoxb7-Cep120 KO and control kidneys. (D) Compbio analysis of pathway networks of the differentially expressed genes upon centrosome loss. (E) Schematic showing KEGG pathway analysis highlighting the Wnt signaling pathway network implicated upon centrosome loss. (F) Differential expression of Wnt family members in kidney cells of Hoxb7-Cep120 KO mice compared to control.

Other pathways impacted upon CL include regulators of the cell apical junction and surface (Figure 4C), indicative of potential defects in apicobasal and/or basolateral polarity during tubule formation ^51-53^. Finally, there was dysregulation of components of G_2_/M progression and p53 signaling (Figure 4C). These pathways are known to be disrupted upon centrosome loss, resulting in activation of the mitotic surveillance pathway leading to cell cycle arrest and p53-mediated apoptosis ^24,25,27,28,39^, consistent with our immunofluorescence analyses (Figures S2C-2F). We also analyzed the 618 DEGs using the CompBio (COmprehensive Multi-omics Platform for Biological InterpretatiOn) tool, which provides a holistic, contextual map of the core biological concepts and themes associated with those pathways ^54,55^. Upon centrosome loss in the developing E13.5 kidney, the super clusters related to mitotic spindle pole, cilium assembly, contractile actin filament, adhesion junctions, cytoskeletal organization, DNA damage, ECM remodeling were the most affected pathways (Figure 4D).

Finally, we performed transcriptional profiling of Pkd1-KO embryonic kidneys at the same embryonic stage, to provide a comparison between CL and ciliary signaling dysfunction. UB progenitors were isolated from Pkd1-KO and WT control at E13.5 by FACS (Figure 4A), and analyzed using bulk RNA sequencing. Surprisingly, the analysis showed no significant DEGs that passed both the >2 log fold-change and p-value < 0.05 thresholds in Pkd1-KO embryonic kidneys compared with WT control at E13.5 (Figure S3A). This suggests that Pkd1 loss has no significant observable transcriptional changes at this early stage, unlike the ablation of centrosomes in the Cep120 knockout mice. Altogether, these data highlight the cellular and molecular pathways that are impacted upon centrosome loss during embryonic kidney development, and which likely contribute to the observed developmental defects.

### Centrosome loss alters Wnt7 and Wnt11 expression patterns during UB branching

Several of the pathways and processes identified in the Hallmark and CompBio analyses are downstream effectors of Wnt signaling (Figure 4E). The impacted downstream effectors of canonical Wnt-β-catenin signaling were regulators of adhesion junctions, TGFβ, MAPK pathways, and cell cycle progression (Figure 4E). This indicates that defective canonical Wnt-β-catenin signaling upon centrosome loss may underlie the abnormal growth and renewal of nephron progenitor cells that occurs during kidney development. In addition, there were changes in downstream effectors of non-canonical Wnt/PCP pathways (Figure 4E), which are essential for cellular motility ^56^, adhesion ^57^, and rearrangements of the cytoskeleton ^58^.

We found that the expression of several Wnt genes were altered in the centrosome-loss group (Figure 4F), some of which (e.g., Wnt 5, 7 and 11) function through the non-canonical Wnt/PCP pathways ^58^. Wnt11 is involved in UB branching morphogenesis and convergent extension during tubulogenesis, and its absence leads to kidney hyperplasia due to defective UB branching ^59^. Wnt11 expression is highly restricted to ureteric bud tip progenitor cells, where it plays an important role in promoting their self-renewal, as well as communication with the cap mesenchymal cells ^59^. Immunofluorescence staining of wildtype kidneys at E13.5 showed that Wnt11 expression was restricted to the ureteric bud tip cells, but not in differentiated stalk cells, as expected (Figures 5A and 5C). However, the expression of Wnt11 was reduced in the UB tip cells in both Six2-Cep120 and Hoxb7-Cep120 KO kidneys, consistent with the RNAseq results (Figures 4F, 5A and 5C).

**Figure 5.**
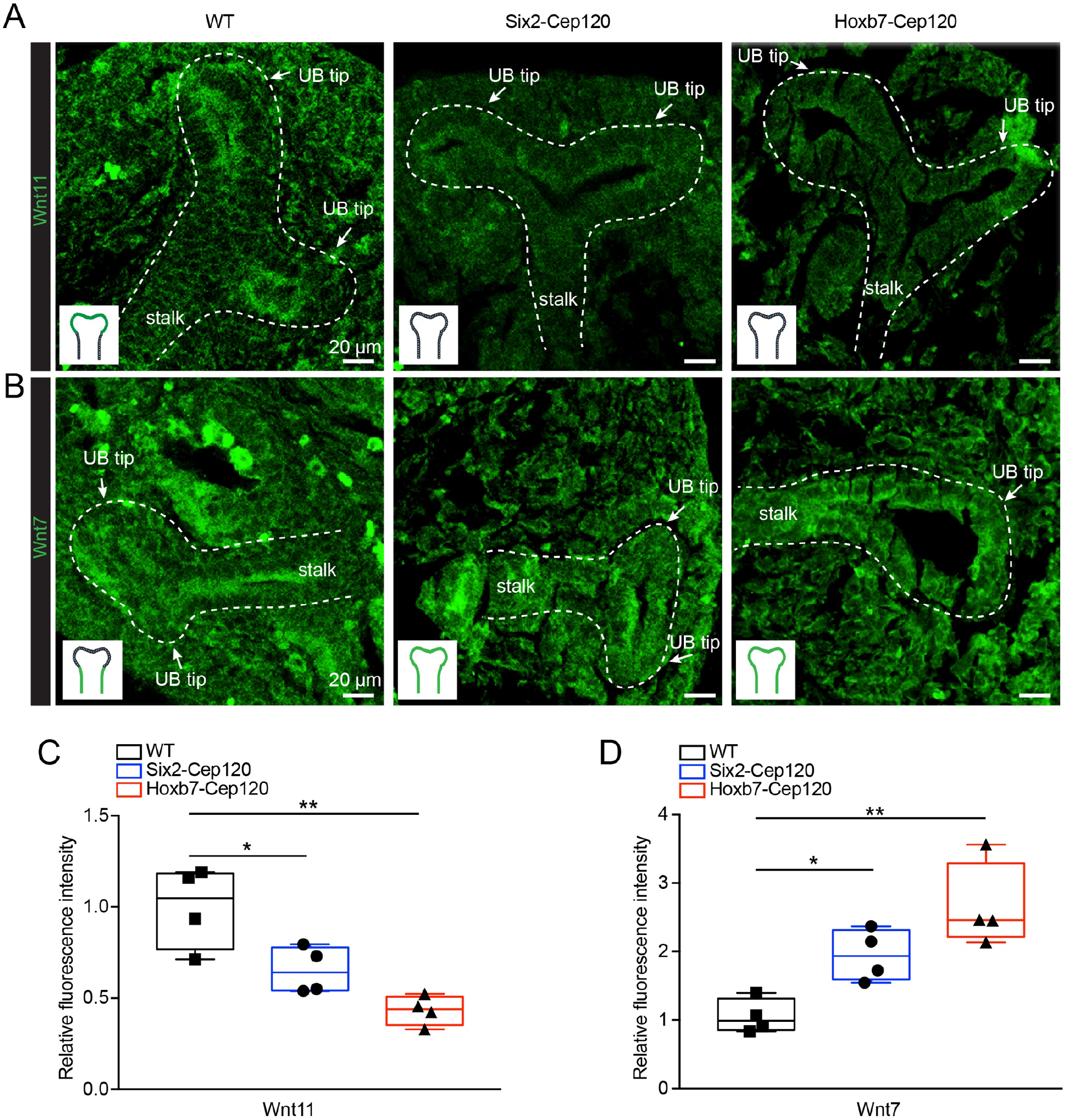
Centrosome loss alters Wnt7 and Wnt11 expression patterns during UB branching. (A) Immunostaining of Wnt11 in kidney sections of Six2-Cep120, Hoxb7-Cep120 and control mice at E13.5. (B) Immunostaining of Wnt7 in kidney sections of E13.5 Six2-Cep120, Hoxb7-Cep120 and control mice. (C) Relative fluorescence intensity measurements of Wnt11 in UB tips and stalks in the nephrogenic zone in kidney sections of Six2-Cep120, Hoxb7-Cep120 and control mice at E13.5. (D) Relative fluorescence intensity measurements of Wnt7 in UB tips and stalks in the nephrogenic zone in kidney sections of Six2-Cep120, Hoxb7-Cep120 and control mice at E13.5. N = 4 mice per group.

In wildtype developing kidneys, Wnt7b expression is restricted to the differentiated ureteric stalk epithelium where it activates Wnt/PCP signaling in the surrounding medullary interstitium ^60^. Wnt7b is required for elongation of the collecting duct and loops of Henle, by controlling the tubular luminal diameter and length. Wnt7b KO kidneys lack normal tubular elongation in the medullary region, suggesting that Wnt7b may act locally in the UB trunk or non-autonomously on the cells adjacent to developing tubule to stabilize polarity of proliferating UB cells ^58^. Immunofluorescence staining of wildtype kidneys at E13.5 showed that Wnt7 expression was exclusively in stalk but not the UB tip cells. In contrast, the expression of Wnt7 was significantly increased in the UB tip cells in both Six2-Cep120 and Hoxb7-Cep120 KO kidneys (Figures 5B and 5D). Altogether, these data suggest that centrosome loss leads to abnormal expression and localization of Wnt7 and Wnt11, resulting in pre-mature differentiation of UB progenitors and abnormal branching morphogenesis.

### Centrosome loss causes early-onset fibrosis and cystogenesis

We next determined the consequences of centrosome loss in postnatal mice. Both Six2-Cep120 and Hoxb7-Cep120 mice were born in the Mendelian ratios (Table 1). Analysis of kidneys isolated from Six2-Cep120 and Hoxb7-Cep120 KO animals at P15 showed a reduction in size compared with wildtype, which was already evident at P0 and became more prominent by P15 (Figures 6A and 6C). Notably, the Cep120 KO kidneys were smaller than Hoxb7-Pkd1 or Six2-Pkd1 KO (Figures 6A and 6C). Importantly, centrosome loss resulted in early onset cyst formation in both KO models (Figure 6A). Quantification of cyst index indicated that cystogenesis in Six2-Cep120 and Hoxb7-Cep120 KO kidneys was milder than that of Six2-Pkd1 and Hoxb7-Pkd1 mutants (Figure 6A and D). To identify the origin of cysts in the knockout kidneys, we stained P15 sections for lotus tetragonolobus lectin (LTL, proximal tubules marker) and dolichos biflorus agglutinin (DBA, collecting ducts marker). In the kidneys of the Six2-Cep120 KO mice ∼66% of the cysts were LTL positive and thus proximal tubule-derived, whereas in Hoxb7-Cep120 KO kidneys approximately ∼75% of cysts were DBA positive (Figure S4A). These results indicate that the majority of cysts are specifically derived from progenitor cells ablated of centrosomes. Indeed, quantification of Cep120, centrosome and cilia number showed a significant reduction in cystic epithelial cells of Cep120-KO mice at P15 (Figures S4B-S4E).

**Figure 6.**
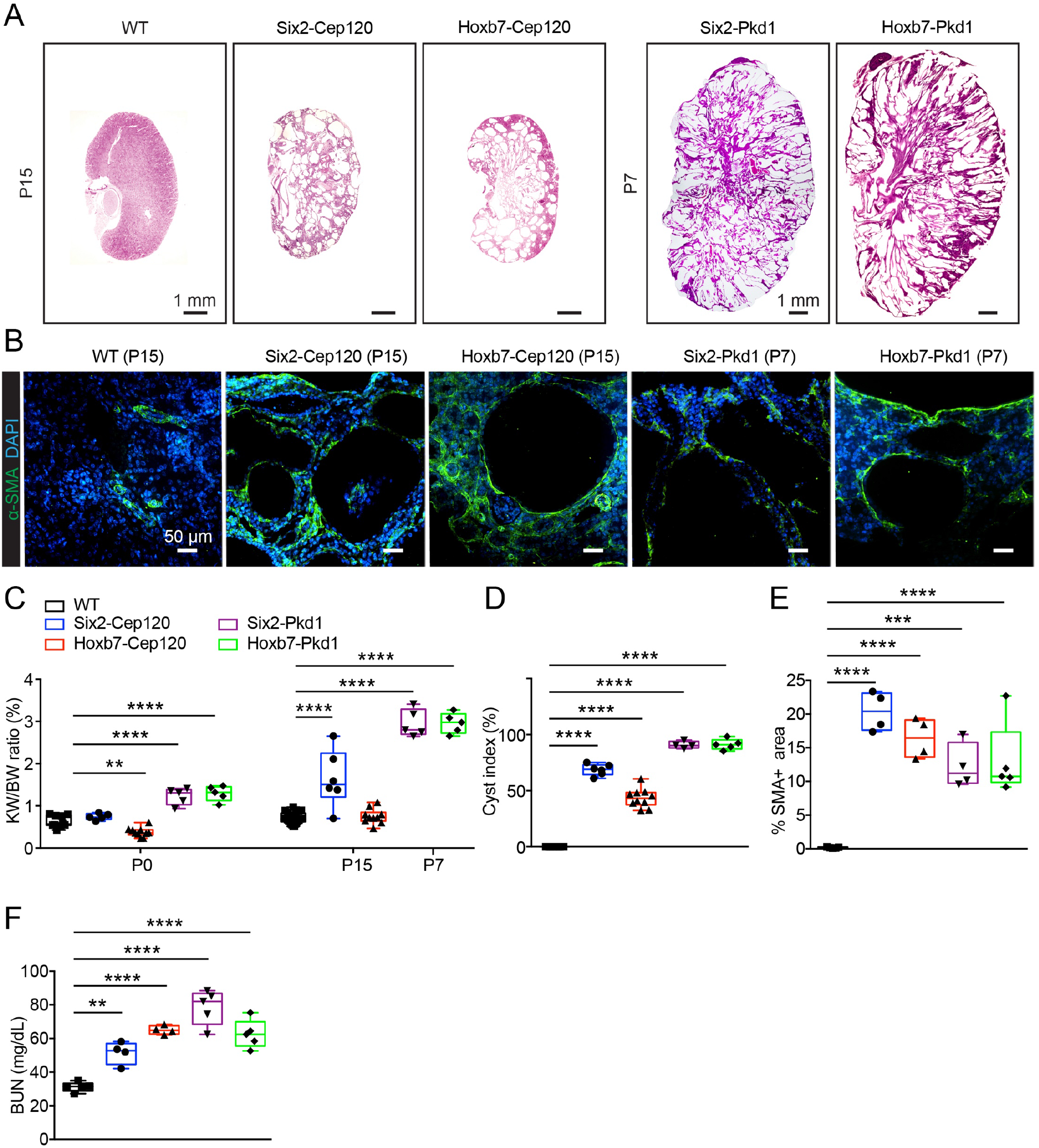
Centrosome loss causes enhanced fibrosis and milder cyst formation when compared to Pkd1 knockout models. (A) H&E staining of postnatal kidney sections of Six2-Cep120, Hoxb7-Cep120, Six2-Pkd1, Hoxb7-Pkd1 and control mice. (B) Immunostaining for fibrosis marker α-SMA in kidney sections of Six2-Cep120, Hoxb7-Cep120, Six2-Pkd1, Hoxb7-Pkd1 and control mice. (C) Kidney-weight-to-body-weight (KW/BW) ratio analysis of kidneys from Six2-Cep120, Hoxb7-Cep120, Six2-Pkd1, Hoxb7-Pkd1 and control mice. N = 4-10 mice per group. (D) Quantification of cystic index of kidneys from Six2-Cep120, Hoxb7-Cep120, Six2-Pkd1, Hoxb7-Pkd1 and control mice. N = 4-10 mice per group. (E) Quantification of α-SMA+ area in kidney sections from Six2-Cep120, Hoxb7-Cep120, Six2-Pkd1, Hoxb7-Pkd1 and control mice. N = 4-5 mice per group (F) Measurement of Blood Urea Nitrogen (BUN) from Six2-Cep120, Hoxb7-Cep120, Six2-Pkd1, Hoxb7-Pkd1 and control mice. N = 4-5 mice per group.

**Table 1:** Number of animals of each genotype isolated from the respective crosses (percentage of the total is indicated in parentheses)

|  | **Cep120F/F**<br>**+/+** | **Cep120F/+**<br>**+/+** | **Cep120F/+**<br>**Hoxb7-Cre/+** | **Cep120F/F**<br>**Hoxb7-Cre/+** | **Total:** |
| --- | --- | --- | --- | --- | --- |
| **P0** | 7 (25%) | 5 (18%) | 8 (29%) | 8 (29%) | 28 |
| **P15** | 8 (32%) | 7 (28%) | 5 (20%) | 5 (20%) | 25 |

|  | **Cep120F/F**<br>**+/+** | **Cep120F/+**<br>**+/+** | **Cep120F/+**<br>**Six2-Cre/+** | **Cep120F/F**<br>**Six2-Cre/+** | **Total:** |
| --- | --- | --- | --- | --- | --- |
| **P0** | 3 (21%) | 3 (21%) | 3 (21%) | 5 (36%) | 14 |
| **P15** | 10 (37%) | 5 (19%) | 7 (26%) | 5 (19%) | 27 |

Next, we investigated the extent of fibrosis in the KO animals. Immunostaining of kidneys isolated at P15 with the fibrotic marker α-SMA showed increased levels in Six2-Cep120 and Hoxb7-Cep120 KO kidneys compared to wildtype (Figures 6B and 6E). The extent of fibrosis was significantly higher than that observed in Hoxb7-Pkd1 or Six2-Pkd1 KO animals (Figures 6B and 6E), suggesting centrosome loss causes more fibrosis compared to Pkd1-loss. Finally, analysis of blood urea nitrogen (BUN) levels showed an increase in Six2-Cep120 and Hoxb7-Cep120 KO mice, but lower than those of Six2-Pkd1 and Hoxb7-Pkd1 KO (Figure 6F). In sum, loss of centrosomes results in small and fibrocystic kidneys after birth that are distinct from those observed in early-onset ADPKD mice.

### Transcriptional profiling of Cep120 fibrocystic kidneys implicates EMT signaling

We sought to identify the defective pathways upon centrosome loss that lead to the cystogenesis and fibrosis phenotypes after birth. Kidneys were collected from Hoxb7-Cep120 KO and WT mice at P15, total RNA extracted and subjected to bulk RNA-seq analysis. There was a total of 94 DEG in Cep120 KO kidneys, including 36 upregulated genes and 58 downregulated genes (Figure 7A). The low number of DEG at this timepoint compared with the embryonic kidneys (Figure 4) is likely because centrosome-less cells were not enriched by FACS, as the Hoxb7-CreGFP expression is embryonic only. Hallmark pathway analysis identified downregulated pathways involved in xenobiotic metabolism and oxidative phosphorylation, which point to metabolic defects and impaired ATP production, all of which can contribute to the development of cystic, fibrotic and chronic kidney disease ^61^. The upregulated gene sets were mainly related to EMT-like signaling (Figure 7B). Previous studies have demonstrated that increased activity of the EMT-like pathway in tubular epithelial cells is associated with enhanced deposition of extracellular matrix, abnormal energy metabolism, and aberrant proliferation of cyst epithelial cells ^62,63^. Similarly, CompBio analysis showed that the most affected signaling networks are ones involved in regulation of interstitial fibroblast differentiation, Wnt/PCP pathway in cell differentiation, and stromal collagen deposition, highlighting the processes that are contributing to the cystic and fibrotic phenotypes upon centrosome loss. (Figure 7C). Of note, we found that Cxcl5 is highly upregulated in our dataset. Cxcl5 signaling via its receptor, Cxcr2, can promote proliferation and metastasis of different cancer cell types and contribute to the EMT of nasopharyngeal carcinoma cells by activating ERK/GSK3β/snail signaling ^64-66^. Altogether, our data suggested that tubular epithelial cells may upregulate Cxcl5-Cxcr2 mediated signaling to promote cystogenesis and fibrosis (Figure 7D).

**Figure 7.**
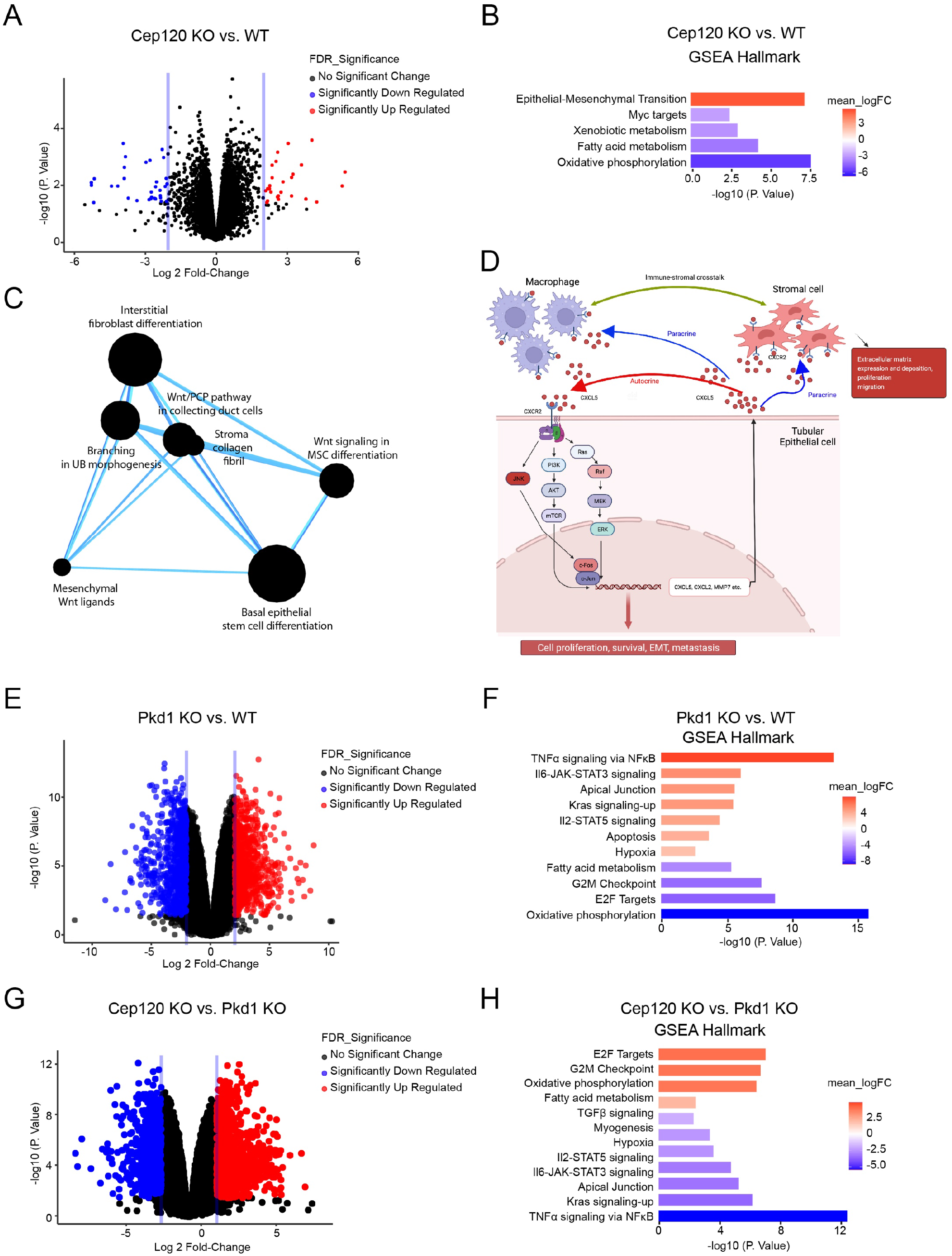
Bulk RNAseq analysis of kidneys from control, Hoxb7-Cep120 and Hoxb7-Pkd1 mice at P15. (A) Volcano plot showing differentially expressed genes in kidneys of Hoxb7-Cep120 compared with control mice. N = 3 mice per group. (B) Hallmark pathway analysis of Hoxb7-Cep120 versus control kidneys. (C) Compbio analysis of pathway networks of the differentially expressed genes upon centrosome loss. (D) Model of Cxcl5-Cxcr2 signaling which may drive proliferation of tubular epithelia and interstitial cells leading to the fibrocystic kidney phenotype upon centrosome loss. (E) Volcano plot showing differentially expressed genes in kidneys of Hoxb7-Pkd1 compared with control. N = 3 mice per group. (F) Hallmark pathway analysis of Hoxb7-Pkd1 versus control kidneys. (G) Volcano plot showing differentially expressed genes in Hoxb7-Cep120 versus Hoxb7-Pkd1 kidneys at P15. N = 3 mice per group. (H) Hallmark pathway analysis of Hoxb7-Cep120 versus Hoxb7-Pkd1 kidneys.

Next, we sought to compare the signaling pathways that are disrupted in cystic kidneys following centrosome loss (Cep120-KO) versus ciliary signaling (Pkd1-KO). To address this, we performed bulk RNA-seq analysis of kidneys isolated from Hoxb7-Pkd1 KO mice at P15. There was a total of 1,550 DEGs including 786 upregulated and 764 downregulated genes in Pkd1-KO kidneys compared with wildtype (Figure 7E). Hallmark analysis identified several signaling pathways known to be involved in cystogenesis and fibrosis in ADPKD (Figure 7F), including TNFα-NF-κB, TGFβ, JAK-STAT, and KRAS signaling ^67,68^. Subsequently, we compared the DEGs from the Cep120-KO and Pkd1-KO mice at P15 (Figure 7G). There was a total of 1,585 DEGs including 1032 upregulated and 553 downregulated genes in the Cep120 KO kidneys compared with Pkd1 (Figure 7G). Hallmark analysis showed that there are significant differences between the two cystic disease models with regards to the cellular processes and pathways impacted (Figure 7H). Unlike Cep120 loss (Figure 7A), the majority of the observed changes in gene expression are derived from ADPKD-deficient kidneys (Figure 7H). The upregulated gene sets in Cep120-KO compared to Pkd1-KO mice include those involved in the G_2_M checkpoint and E2F targets, which indicate that centrosome loss causes more severe cell cycle defects compared ADPKD models (Figure 7H). Compared with Pkd1 loss, many signaling pathways involved in cystogenesis in ADPKD such as TNFα-NF-κB, TGFβ, JAK-STAT, and KRAS signaling, as well as the inflammatory response are significantly lower in Cep120 KO kidneys. This suggests that these signaling pathways are less activate, and may explain the reduced severity of the cystic phenotype in Cep120 relative to Pkd1 KO kidneys.

## Discussion

In this study, we examined the consequences of disrupting centrosome biogenesis in nephron progenitor cells during embryonic kidney development. We observed that kidneys formed abnormally, were dysplastic and small in size, and became rapidly fibrotic and cystic postnatally. Ablation of Cep120 disrupted the balance in the growth and differentiation of nephron progenitor cells, resulting in depletion of both the MM and UB pool of progenitors. This led to premature cell differentiation and defective branching morphogenesis, ultimately leading to overall reduced nephron formation and abundance. The congenital developmental defects, as well as the early-onset fibrocystic phenotypes, in the Cep120-KO mice recapitulate the pathological kidney characteristics observed in NPH, JS and JATD ciliopathy patients. Importantly, the observed changes in progenitor cell growth and fate, nephron development, and kidney morphology were different from the conditional ablation of *PKD1* in the same progenitor niches. Although the deletion of *PKD1* in mice causes a severe early-onset cystic disease phenotype that does not mimic the slow, degenerative human disease pathology ^69^, we ablated the gene in the same progenitor cells to compare the consequences of defective centrosome biogenesis to ciliary signaling. Our results indicate that mutations which disrupt centrosome biogenesis and function cause defects in kidney progenitor cells that are more pronounced than defective ciliary signaling. Indeed, mutations in several other genes involved in centrosome biogenesis, such as POC1B and C2CD3, cause ciliopathies like JS and JATD, and these patients develop the more severe early-onset fibrocystic kidney disease ^70,71^.

We wondered why loss of Cep120 and centrosomes resulted in reduced abundance of nephron progenitor cells, and development of the small, dysplastic kidney phenotype. Defective centrosome biogenesis can result in the loss of progenitor cells by a combination of cell cycle arrest, cell death, or premature differentiation ^27,28^. We found that induction of CL in both MM and UB progenitors resulted in prolonged mitosis, nuclear enrichment of p53, and caspase-mediated apoptosis (Figures S2A-S2F). This is consistent with previous studies where depletion of centrosome proteins in neural progenitor cells in the brain causes prolonged mitosis, and activation of the p53-mediated mitotic surveillance pathway, resulting in a reduced progenitor pool and differentiated neurons ^27^. In contrast, loss of centrosomes in Sox9-positive lung progenitors, or in endodermal cells of the developing intestines, does not induce p53-mediated apoptosis. It has been proposed that high expression and activity of the ERK signaling pathway in the lung and intestinal progenitor cells confers protection against the p53-mediated apoptosis^28^. However, this protective mechanism is unlikely to be in play in kidney progenitors, since there is consistent expression and activity of ERK in the MM and UB progenitor cells ^72,73^, yet they do not survive following centrosome loss. Moreover, it is unclear why, following the differentiation of the MM and UB into tubular epithelia, cells can survive and proliferate in the absence of centrosomes.

One possibility is that components of the mitotic surveillance pathway are differentially expressed in the nephron progenitor stage compared to differentiated tubular cells, such that the pathway is only activated in progenitors. Indeed, our RNAseq analyses indicate that activators of the p53-mediated surveillance pathway (e.g., Pidd1, ^74,75^) were highly upregulated upon centrosome loss during the progenitor cell stage (E13.5, Figure 4), but were not differentially expressed in fully differentiated tubular cells at P15. Conversely, the expression of inhibitors of the p53-mediated pathway (e.g., Mdm2, ^76^) were downregulated in centrosome-less cells during embryonic stages, but the expression was high in postnatal differentiated epithelial cells (Supplementary Excel 2). Thus, we believe that the observed apoptosis of the renal progenitor population may be at least partially a result of the p53-mediated surveillance mechanism. One way to test this would be to delete one or both copies of p53, which can enhance survival of cells lacking centrosomes in certain tissues ^26-28,39^. However, conditional inactivation of p53 signaling in embryonic kidneys delays UB branching, disrupts proliferation of MM, and causes renal hypoplasia ^77,78^. Thus, we could not test the effects of blocking the p53-mediated surveillance pathway in centrosome-less kidneys directly.

Another possibility for why loss of centrosomes disrupted nephron development and led to the dysplastic kidney phenotype is likely due to abnormal differentiation of the progenitor cells. Prior studies have established an important role for centrosomes in stem cell renewal and differentiation in various organs and organisms. In the developing *Drosophila* brain, the neuron stem cells undergo multiple rounds of asymmetric cell division to generate a stem cell and a differentiating neuron ^79^. Ablating centrosomes in the neural stem cells disrupts the balance of asymmetric divisions, leading to increased death or premature differentiation of the progeny from these divisions ^80,81^, and ultimately depletion of the progenitor pool. Similarly, disrupting centrosome biogenesis and inheritance in the mouse brain results in abnormal asymmetric stem cell division, depletion of progenitors from the ventricular zone, leading to the small brain phenotype ^27^. The premature differentiation of progenitor cells is thought to occur due to abnormal inheritance of the older versus younger centrosome in an asymmetrically dividing cell ^82,83^. This asymmetric centrosome inheritance is linked to the segregation of fate determinants. For example, fate-determining mRNAs are associated with one centrosome during cell divisions of the mollusc embryo, governing binary fate decision ^84^, while a regulator of Notch ligand activity, Mindbomb1, was found to localize asymmetrically to the daughter centrosomes in chick neural progenitors, leading to its segregation to prospective neurons during mitosis ^85^. Another intriguing possibility by which centrosome loss may contribute to cell fate decisions is through a differential ability of the dividing daughter cells to assemble primary cilia. Cells that inherit the older centrosome assemble cilia earlier, and respond to Hedgehog signaling, which promotes stem cell identity. On the other hand, their siblings that inherit the younger centrosome do not self-renew due to lower Hedgehog activity and thus commit to differentiation ^86,87^. In the kidney, our data indicate that CL in the renal progenitors leads to premature cell differentiation of both the MM and UB niches, and accelerated formation of pretubular aggregates. Of note, the RNAseq analysis of Cep120-KO embryonic kidneys identified changes in the Wnt signaling pathway upon centrosome loss. Components of Wnt signaling have been shown to localize to centrosomes, and to regulate cell fate. In C. elegans, centrosomal localization of the β-catenin homologue SYS-1 promotes dynamic centrosome–associated proteasomal degradation during asymmetric cell division, limiting its retention by daughter cells prior to cell fate specification. Dishevelled (Dvl) mediates Wnt signaling that is known to be critical for cell-fate determination and planar cell polarity ^88^. Diversin, a known antagonist of β-catenin stability, depends on centrosomal localization to appropriately promote β-catenin degradation ^89^. Cells without centrosomes undergo attenuated response to Wnt and accumulate a distinct, higher-molecular-weight species of pβ-catenin which results in cell fate changes ^88,89^. In sum, we interpret our result to indicate that the reduced abundance of progenitor cells and nephrons observed upon CL is likely a combination of cell cycle delay, cell death, and Wnt-dependent premature differentiation.

In contrast to the progenitor cells, loss of Cep120 and centrosomes led to rapid epithelial cell growth and cyst formation after birth. This suggests that, unlike the brain and lung, CL in the kidney shows differential effects in the differentiated population of cells. Differentiated kidney cells may be more resistant to CL, or protected against the cell cycle arrest and cell death observed in the progenitor stages. One possibility is that the pro-inflammatory and pro-fibrogenic signaling pathways that are elevated upon CL help protect the cells against CL and permit their growth. Our RNAseq analyses of postnatal Cep120-KO fibrocystic kidneys identified a small group of DEGs with strong signatures of EMT and pro-inflammatory signaling, suggesting these pathways may be key for cell survival and growth in the absence of centrosomes, and progression to the fibrocystic kidney phenotype. Among the DEGs, Cxcl5 is the 9^th^ highest upregulated gene (Supplementary Excel 2). The Cxcl5-Cxcr2 signaling axis can promote proliferation and metastasis of different cancer cell types via autocrine signaling and participate in the recruitment of immune cells and promote angiogenesis, tumor growth, and metastasis via paracrine signaling ^64,65^. Analysis of existing single nuclear RNAseq of injured mouse kidneys indicates that Cxcl5 is highly expressed in different segments of nephron epithelial cells, while Cxcr2 is highly expressed in epithelial cells, immune cells and interstitial cells^90,91^. Inhibition of Cxcr2 protects against tubular cell senescence and renal fibrosis following unilateral ureteral obstruction (UUO) ^92^, and reduces kidney injury associated with a reduction of inflammatory cytokines and neutrophils infiltration ^93^. Intriguingly, it was shown that liver cysts of ADPKD patients secrete agonists of Cxcr2, which promote cell proliferation and cyst growth ^94^. We hypothesized that tubular epithelial cells may upregulate Cxcl5-Cxcr2 mediated signaling to promote cystogenesis and fibrosis upon CL. Our future studies will aim to test inhibitors of the Cxcl5-Cxcr2 axis to block the abnormal proliferation of epithelial cells, as well as the crosstalk between the epithelial, immune, and the interstitial cells, and examine the effects in disease progression.

Compared with Pkd1-KO mice, many signaling pathways involved in cystogenesis in ADPKD are significantly lower in Cep120 KO kidneys (Figure 7H). This suggests that these signaling pathways are less activate in centrosome-less kidneys and may explain the reduced severity of the cystic phenotype in Cep120-KO relative to Pkd1-KO kidneys. In contrast, there were upregulated gene sets in Cep120-KO mice, including G_2_M checkpoint and E2F targets, which indicate that centrosome loss causes more severe cell cycle defects compared with ADPKD models, which may explain the differential effects on the cells and the dysplastic kidney phenotype.

Finally, we wondered how mutations in Cep120 (or other centrosome biogenesis proteins) that manifest in the nephron progenitors contribute to the overall renal dysplasia, fibrosis and cyst formation that occurs in NPH, JS and JATD patients. In the accompanying study, Langner and colleagues (2023) induced conditional deletion of Cep120 in the stromal progenitor cells of the developing kidney. Cep120 deletion disrupted centrosome biogenesis in stromal progenitor-derived cell types including pericytes, interstitial fibroblasts, mesangial, and vascular smooth muscle cells, and resulted in reduced abundance of several stromal cell lineages (interstitial pericytes, interstitial fibroblasts, and mesangial cells). This led to development of small, hypoplastic kidneys with visible signs of medullary atrophy and delayed nephron maturation by postnatal day 15. The reduced interstitial cell populations were due to a combination of defective cell cycle progression of stromal progenitors lacking centrosomes, p53-mediated apoptosis, and changes in signaling pathways essential for differentiation of stromal lineages. This indicates that aberrant centrosome biogenesis in all three kidney progenitor populations is contributing to the small dysplastic/hypoplastic kidney phenotype in patients. Yet, there was no spontaneous fibrosis or early-onset cystogenesis after birth upon CL in the stromal progenitors, unlike the those observed in the Six2-Cep120 and Hoxb7-Cep120 KO mice (Figure 6). However, CL in the interstitium sensitized kidneys of adult mice, causing rapid fibrosis via enhanced TGFβ/Smad3-Gli2 signaling after renal injury (Langner et al, 2023). Collectively, our data indicate that mutations in Cep120 and centrosome biogenesis genes in the nephron progenitors plays a major role in spontaneous fibrosis and cystogenesis, while defects in stromal progenitors contribute to the congenital developmental defects and enhanced fibrosis following renal injury in the patients. In sum, our study provides a detailed characterization of the underlying molecular and cellular defects in renal centrosomopathies ^95,96^.

## Acknowledgments

We would like to thank the Washington University Genome Technology Access Center (GTAC) for help with bioinformatic analyses. We also thank Dr. Jeff Miner for sharing reagents used in this study. We also thank members of the Mahjoub lab and the Washington University Ciliopathy Research Group for helpful feedback on this project, and critical reading of the manuscript. Finally, we thank members and the Washington University Center for Cellular Imaging (WUCCI) for assistance with light sheet microscopy and Imaris data analysis (supported in part by The Children’s Discovery Institute of Washington University and St. Louis Children’s Hospital (CDI-CORE-2015-505 and CDI-CORE-2019-813) and the Foundation for Barnes-Jewish Hospital (3770 and 4642). This study was supported by funding from the National Institute of General Medical Sciences (HHS - NIH) (R01GM140115) to B.W. and the National Institute of Diabetes and Digestive and Kidney Diseases (R01-DK108005) to M.R.M.

## Author contributions

T.C. performed all the experimental work and data quantitation, prepared the figures, and wrote the manuscript. C.A. helped with mouse genotyping and immunostaining experiments. K.S. assisted with mouse mating, genotyping and tissue isolation. B.W. developed and shared the *Cep120^F/F^* mouse model. S.J. helped with RNAseq analysis. M.R.M. conceived the project, helped write the manuscript, and supervised all aspects of the study. All authors reviewed and edited the manuscript.

## Materials and Methods

### Generation of mouse models

All animal studies were performed following protocols that are compliant with guidelines of the Institutional Animal Care and Use Committee at Washington University and the National Institutes of Health. Conditional *CEP120* and *PKD*1 knockout strains were generated by crossing *Cep120^F/F^* or *PKD1^F/F^* mice with *Six2-Cre* or *Hoxb7-Cre-*expressing *strains*, respectively. Litters were genotyped by PCR utilizing primers for *Cep120*: Flox Forward: 5’-CCT CTG CCT CCT TAG TGG ATC-3’; WT Forward: 5’-ATC ACT GTG GAG CCT TGG GCA-3’; Reverse: 5’-TGT TAC TCA GCA GCT GGT ACC-3’; primers for *PKD1*: Forward: 5’-GCC CAC AGC TAT TGT TCC TAA-3’; Reverse: 5’-GGA TAA AGT GAT CAA GCA GCA-3’; primers for Cre: Forward: 5’-CCA ATT TAC TGA CCG TAC ACC-3’; Reverse: 5’-CGT AAC AGG GTG TTA TAA GCA A-3’.

### Histology and Immunofluorescence

Both kidneys were isolated from embryos (E13.5 and E17.5) or postnatal mice (P0 and P15), fixed with 4% paraformaldehyde in PBS for 24 h at 4°C, then embedded in paraffin. Kidneys blocks were sectioned at 5-7 µm thickness using a microtome (Leica; RM2125 RTS), placed onto microscope slides (Thermo Fisher Scientific), and stored at room temperature. Hematoxylin and eosin (H&E) staining of paraffin-embedded sections was performed using standard protocols.

For immunofluorescence staining of paraffin-embedded sections, antigen unmasking was performed by boiling the slides in antigen-retrieval buffer (10 mM Tris Base, 1 mM EDTA, and 0.05% Tween-20, pH 9.0) for 30 min. Samples were permeabilized with 0.05% Triton X-100 in PBS (PBS-T) for 10 min at room temperature, incubated in blocking buffer (3.0% BSA and 0.1% Triton X-100 in PBS) for 1 h, followed by staining with primary antibodies overnight at 4°C (Table 2). After 3 washes with PBS-T, samples were incubated with secondary antibodies for 1 h at room temperature. All Alexa Fluor dye–conjugated secondary antibodies (Thermo Fisher Scientific) were used at a final dilution of 1:500. Nuclei were stained with DAPI, and specimens mounted using Mowiol containing n-propyl gallate (Sigma-Aldrich). Images were captured using a Nikon Eclipse Ti-E inverted confocal microscope equipped with 10x Plan Fluor (0.30 NA), 20x Plan Apo air (0.75 NA), 40x Plan Fluor oil immersion (1.30 NA), 60x Plan Fluor oil immersion (1.4 NA), or 100x Plan Fluor oil immersion (1.45 NA) objectives (Nikon). A series of digital optical sections (z-stacks) were captured using a Hamamatsu ORCA-Fusion Digital CMOS camera at room temperature, and three-dimensional image reconstructions were produced. Images were processed and analyzed using Elements AR 5.21 (Nikon), Adobe Illustrator and Photoshop software.

**Table 2:** List of antibodies used in this study

| **Antibodies (clone #)** | **Product details** | **Dilution** |
| --- | --- | --- |
| Mouse anti γ-tubulin (GTU-88) | Sigma-Aldrich (T5326) | 1:500 |
| Rabbit anti p53 | Leica (NCL-L-p53-CM5p) | 1:50 |
| Mouse anti α-SMA | Sigma-Aldrich (A2547) | 1:500 |
| Rabbit anti phospho-Histone H3 | Cell Signaling (9701) | 1:200 |
| Mouse anti phospho-Histone H3 | Cell Signaling (9706) | 1:200 |
| Rat anti-Cep120 | Produced in-house | 1:500 |
| Rabbit anti cleaved caspase 3 | Cell Signaling (9664) | 1:50 |
| Mouse anti E-Cadherin | BD Bioscience (BDB610181) | 1:500 |
| Rabbit anti Six2 | ProteinTech (11562-1-AP) | 1:200 |
| Rabbit anti Pax2 | BioLegend (901001) | 1:500 |
| Mouse anti WT1 | Abcam (ab93050) | 1:100 |
| Rabbit anti Wnt11 | GeneTex (GTX105971) | 1:50 |
| Rabbit anti Wnt7 | ProteinTech (10605-1-AP) | 1:50 |
| Mouse anti Ki67 | Cell Signalling (9449) | 1:1000 |
| Rabbit anti-podocin | Roselli et al., 2002 | 1:5,000 |

### Quantification of Cep120 and centrosome number

For quantification of both Cep120 expression and centrosome number in kidney epithelial cells, kidney sections were stained with antibodies against Cep120 and γ-tubulin. Normal centrosome number was defined as cells containing one or two foci of γ-tubulin, while loss of centrosomes was defined as cells containing zero foci.

### Quantification of the density metanephric mesenchyme, renal vesicles, comma-shaped bodies and S-shaped bodies structures

To determine the density of Six2-positive metanephric mesenchyme at different developmental stages, sections from wildtype, Six2-Cep120, Hoxb7-Cep120, Six2-Pkd1 and Hoxb7-Pkd1 kidneys at E13.5 and E17.5 were immunostained for Six2 (Table 2). For quantification of the density of the intermediate structures of the development of nephron: renal vesicles (RV), comma-shaped bodies (CSB) and S-shaped bodies (SSB), samples were stained with antibodies against theses structure markers Pax2 and WT1. The relative density of metanephric mesenchyme was determined by quantifying the number of Six2-positive cells per unit area. Similarly, the relative abundance of RV, CSB and SSB structures was measured per unit area. The density of these structures was calculated by dividing the total number of each structure by the whole area of the whole kidney section area. Kidneys of five mice were analyzed per genotype.

### Evaluation of postnatal kidney morphology and function

Kidney weight (KW) and body weight (BW) were measured following isolation of kidneys at P0 and P15. KW/BW ratio was then calculated by dividing the average kidney weight by body weight (grams).

The entire sagittal kidney section was used to calculate cyst burden. Sagittal kidney sections were stained with hematoxylin and eosin (H&E) and examined by light microscopy. All kidneys were imaged under the same magnification. ImageJ analysis software was used to calculate the cyst index, which refers to the cumulative area of cysts within the total area of the kidney.

For quantification of kidney fibrosis, kidney sections from wildtype, Six2-Cep120, Hoxb7-Cep120, Six2-Pkd1 and Hoxb7-Pkd1 mice at P7 and P15 were immunostained with antibodies against α-SMA. Multiple regions of each kidney were imaged using a 20x objective then analyzed using ImageJ software. The data are expressed as the mean area of α-SMA+ foci per unit area (mm^2^) of each kidney section.

For analysis of kidney function, the level of blood urea nitrogen (BUN) was measured in mice at P15. Blood serum was separated (6,000 g, 15 minutes at 4°C) from blood isolated by sub-mandibular bleeding and used for the BUN assay (BUN-Urea, BioAssay Systems) following the manufacturer’s protocol.

### Whole organ clearing, immunostaining and Light Sheet microscopy

Kidneys of wildtype, Six2-Cep120, Hoxb7-Cep120 at E15.5 were isolated, fixed in 4% paraformaldehyde in PBS for 24 h at 4 °C, washed with PBS for three times for 2 h each at 4°C. Whole kidneys were placed in CUBIC clearing buffer (25% Urea, 25% Quadrol (N,N,N′,N′-Tetrakis (2-hydroxypropyl)ethylenediamine; EDTP), 15% Triton X-100 in ddH_2_O) for 3 days with shaking at 37°C. The cleared, transparent kidneys were then washed three times with PBS-T for 2 h each at 4 °C, and stained with antibodies against E-Cadherin (Table 2) for 48 h at 4°C. Subsequently, samples were washed with PBS-T three times for 2 h each at 4 °C, then incubated with secondary antibody (Goat anti-mouse IgG2a Alexa Fluor 594, 1:500) for 48 h at 4°C, washed with PBS-T three times for 2 h each at 4 °C. Samples were then transferred into the 20-30 mL of CUBIC imaging buffer (RI 1.46, Histodenz, (Sigma D2158) in 0.02 M PBS-0.1% Triton X-100, 0.01% sodium azide, pH7.5) in a 50 mL Falcon tube, then imaged on a Zeiss Lightsheet 7 microscope.

### FACS sorting and RNAseq analysis

For analysis of GFP-positive UB cells with and without centrosomes during embryogenesis, kidneys of wildtype and Hoxb7-Cep120 mice were isolated at E13.5 and dissociated into single cell suspension as previously described ^97^. Briefly, E13.5 embryonic kidneys were dissected and collected in 1.5 ml tube. Collected kidneys were suspended with Trypsin-EDTA solution (Sigma, T3924) with 200 μg/ml DNase I (Sigma) and incubated for 10 min at 37 °C. After the incubation the specimens were triturated with a P-200 pipette and rinsed once with DMEM/F12 containing 10% FCS to inactivate Trypsin. The cell preparation was then treated with collagenase dissolved in DMEM/F12 (1 mg/ml) and incubated at 37 °C for 10 min. The specimens were again triturated and fully dissociated cells were rinsed twice in 500 µl of PBS/ 5% FCS and subjected to Fluorescence active cell sorting (FACS). FACS sorting (Sony Synergy-HAPS 1, 100 micron) was performed using the GFP channel.

For analysis of gene expression changes in fibrocystic kidneys of postnatal mice, whole kidneys were isolated from wildtype and Hoxb7-Cep120 animals at P15. The kidneys were weighed and then grounded using pestle and mortar in liquid N_2_ bath and then lysed in Trizol, then total RNA was isolated using Direct-zol^TM^ RNA MiniPrep Plus (Zymo Research).

For bulk RNAseq analysis, samples were prepared according to library kit manufacturer’s protocol, indexed, pooled, and sequenced on an Illumina HiSeq. Basecalls and demultiplexing were performed with Illumina’s bcl2fastq software and a custom python demultiplexing program with a maximum of one mismatch in the indexing read. RNA-seq reads were then aligned to the Ensembl release 76 primary assembly with STAR version 2.5.1a. Gene counts were derived from the number of uniquely aligned unambiguous reads by Subread:featureCount version 1.4.6-p5. Isoform expression of known Ensembl transcripts were estimated with Salmon version 0.8.2. Sequencing performance was assessed for the total number of aligned reads, total number of uniquely aligned reads, and features detected. The ribosomal fraction, known junction saturation, and read distribution over known gene models were quantified with RSeQC version 2.6.2.

All gene counts were then imported into the R/Bioconductor package EdgeR and TMM normalization size factors were calculated to adjust for samples for differences in library size. Ribosomal genes and genes not expressed in the smallest group size minus one samples greater than one count-per-million were excluded from further analysis. The TMM size factors and the matrix of counts were then imported into the R/Bioconductor package Limma. Weighted likelihoods based on the observed mean-variance relationship of every gene and sample were then calculated for all samples with the voomWithQualityWeights. The performance of all genes was assessed with plots of the residual standard deviation of every gene to their average log-count with a robustly fitted trend line of the residuals. Differential expression analysis was then performed to analyze for differences between conditions and the results were filtered for only those genes with Benjamini-Hochberg false-discovery rate adjusted p-values less than or equal to 0.05.

For each contrast extracted with Limma, global perturbations in known Gene Ontology (GO) terms, MSigDb, and KEGG pathways were detected using the R/Bioconductor package GAGE to test for changes in expression of the reported log 2 fold-changes reported by Limma in each term versus the background log 2 fold-changes of all genes found outside the respective term. The R/Bioconductor package heatmap3 was used to display heatmaps across groups of samples for each GO or MSigDb term with a Benjamini-Hochberg false-discovery rate adjusted p-value less than or equal to 0.05. Perturbed KEGG pathways where the observed log 2 fold-changes of genes within the term were significantly perturbed in a single-direction versus background or in any direction compared to other genes within a given term with p-values less than or equal to 0.05 were rendered as nnotated KEGG graphs with the R/Bioconductor package Pathview.

To find the most critical genes, the raw counts were variance stabilized with the R/Bioconductor package DESeq2 and was then analyzed via weighted gene correlation network analysis with the R/Bioconductor package WGCNA. Briefly, all genes were correlated across each other by Pearson correlations and clustered by expression similarity into unsigned modules using a power threshold empirically determined from the data. An eigengene was then created for each de novo cluster and its expression profile was then correlated across all coefficients of the model matrix. Because these clusters of genes were created by expression profile rather than known functional similarity, the clustered modules were given the names of random colors where grey is the only module that has any pre-existing definition of containing genes that do not cluster well with others. These de-novo clustered genes were then tested for functional enrichment of known GO terms with hypergeometric tests available in the R/Bioconductor package clusterProfiler. Significant terms with Benjamini-Hochberg adjusted p-values less than 0.05 were then collapsed by similarity into clusterProfiler category network plots to display the most significant terms for each module of hub genes in order to interpolate the function of each significant module. The information for all clustered genes for each module were then combined with their respective statistical significance results from Limma to determine whether or not those features were also found to be significantly differentially expressed.

### CompBio analysis

The Differentially Expressed Genes (DEGs) identified by bulk RNAseq at E13.5 and P15 were further analyzed using the CompBio (COmprehensive Multi-omics Platform for Biological InterpretatiOn) software package. CompBio uses contextual language processing and a biological language dictionary that is not restricted to fixed pathway and ontology knowledge bases, such as KEGG or Gene Ontology, to extract “knowledge” from all PubMed abstracts that reference entities of interest. Conditional probability analysis calculated the statistical enrichment of biological concepts (processes/pathways) over those that occur by random sampling. Related concepts built from the list of differentially expressed entities were further clustered into higher-level themes (e.g., biological pathways/processes, cell types and structures, etc.). Themes generated by CompBio incorporate genes in each pathway or process, independent of whether they have a positive or negative function.

### Statistical analyses

Statistical analyses were performed using GraphPad PRISM 9.0. The vertical segments in box plots show the first quartile, median, and third quartile. The whiskers on both ends represent the maximum and minimum values for each dataset analyzed. Collected data were examined by ANOVA or two-tail unpaired t-test as specified in the figure legends. Statistical significance was set at p< 0.05 (*), p<0.01 (**); p<0.001 (***); p<0.0001 (****).

---

**Supplementary Figure 1.**
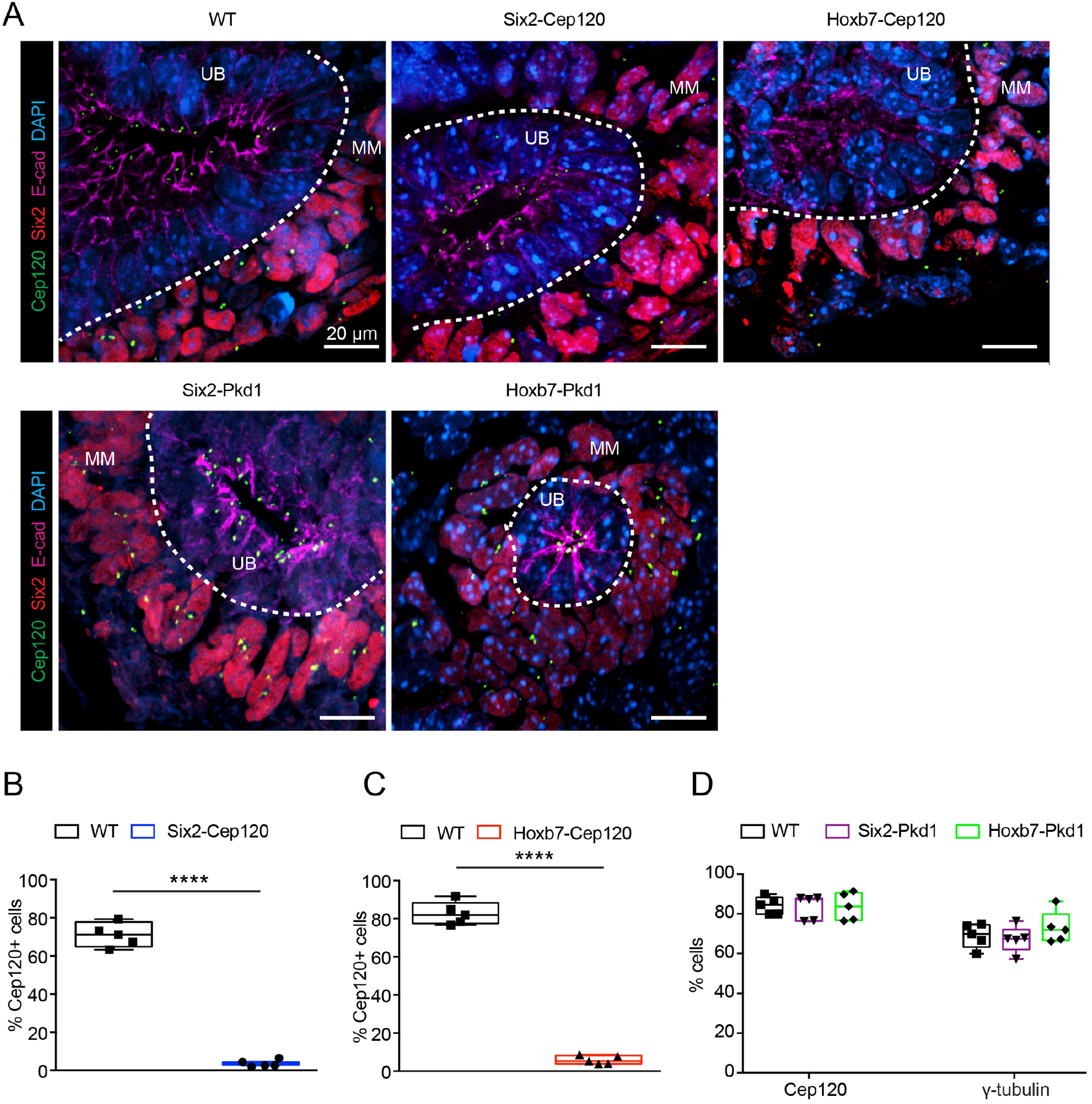
Conditional depletion of Cep120 in nephron progenitors. (A) Immunostaining for Cep120, MM (Six2) and UB (E-cadherin) in kidney sections of Six2-Cep120, Hoxb7-Cep120, Six2-Pkd1, Hoxb7-Pkd1 KO and control mice. (B) Quantification of the percentage of cells lacking Cep120 in MM population of Six2-Cep120 KO and control kidneys. (C) Quantification of the percentage of cells lacking Cep120 in UB population of Hoxb7-Cep120 KO and control kidneys. (D) Quantification of the percentage of cells lacking Cep120 and γ-tubulin of Six2-Pkd1, Hoxb7-Pkd1 KO and control kidneys. N = 5 mice per group.

**Supplementary Figure 2.**
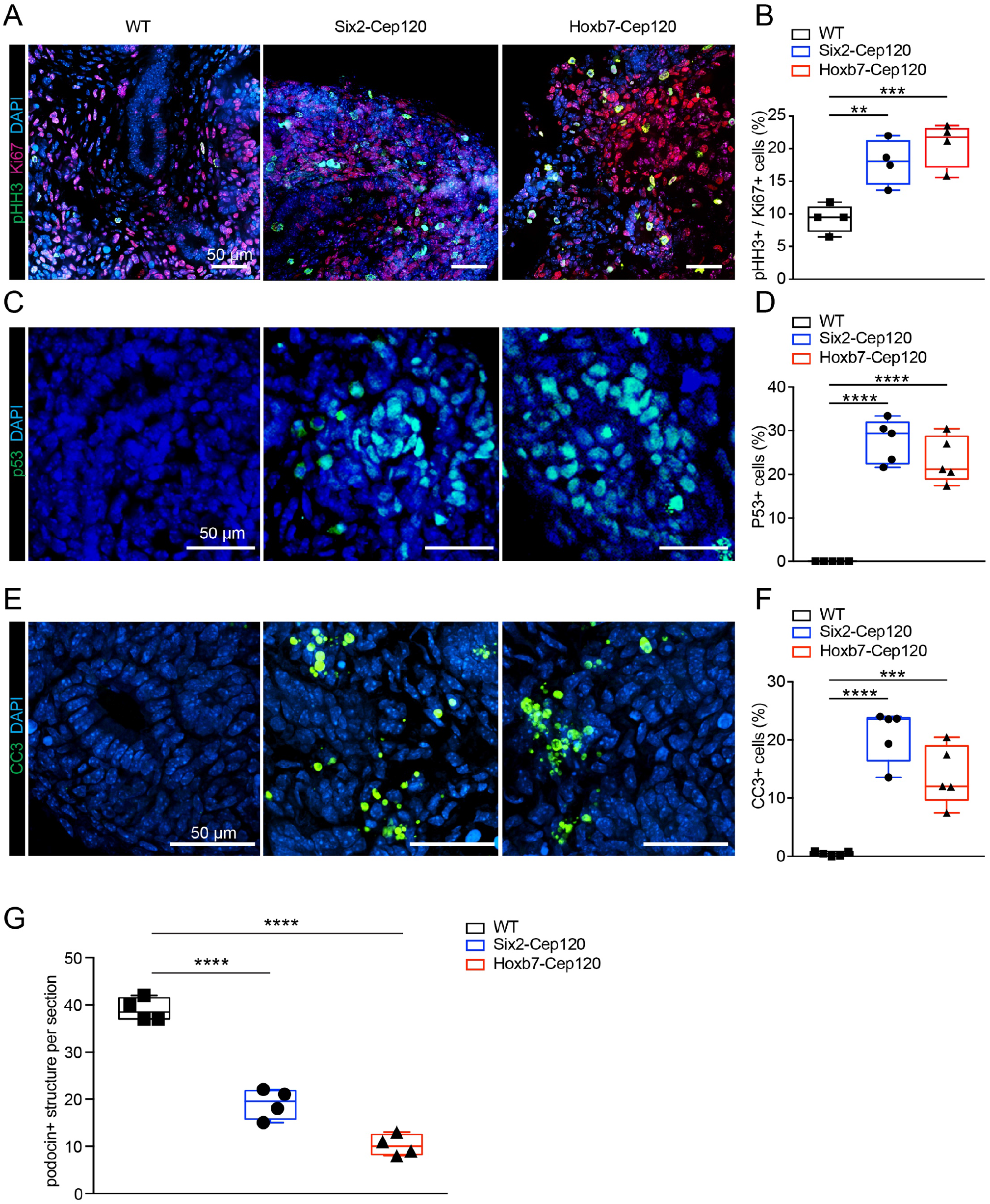
Centrosome Loss activate the mitotic surveillance pathway. (A) Immunostaining for pHH3 and Ki67 in kidney sections of Six2-Cep120, Hoxb7-Cep120 KO and control mice. (B) Mitotic index analysis of Six2-Cep120 KO and control kidneys. N = 4 mice per group. (C) Immunostaining for p53 in kidney sections of Six2-Cep120, Hoxb7-Cep120 KO and control mice. (D) Quantification of cells with nuclear-located p53 of Six2-Cep120, Hoxb7-Cep120 KO and control kidneys. N = 5 mice per group. (E) Immunostaining for Cleaved Caspase 3 in kidney sections of Six2-Cep120, Hoxb7-Cep120 KO and control mice. (F) Quantification of CC3+ cells of Six2-Cep120, Hoxb7-Cep120 KO and control kidneys. N = 5 mice per group. (G) Quantification of podocin^+^ structures of Six2-Cep120, Hoxb7-Cep120 KO and control kidneys at P0. N = 4 mice per group.

**Supplementary Figure 3.**
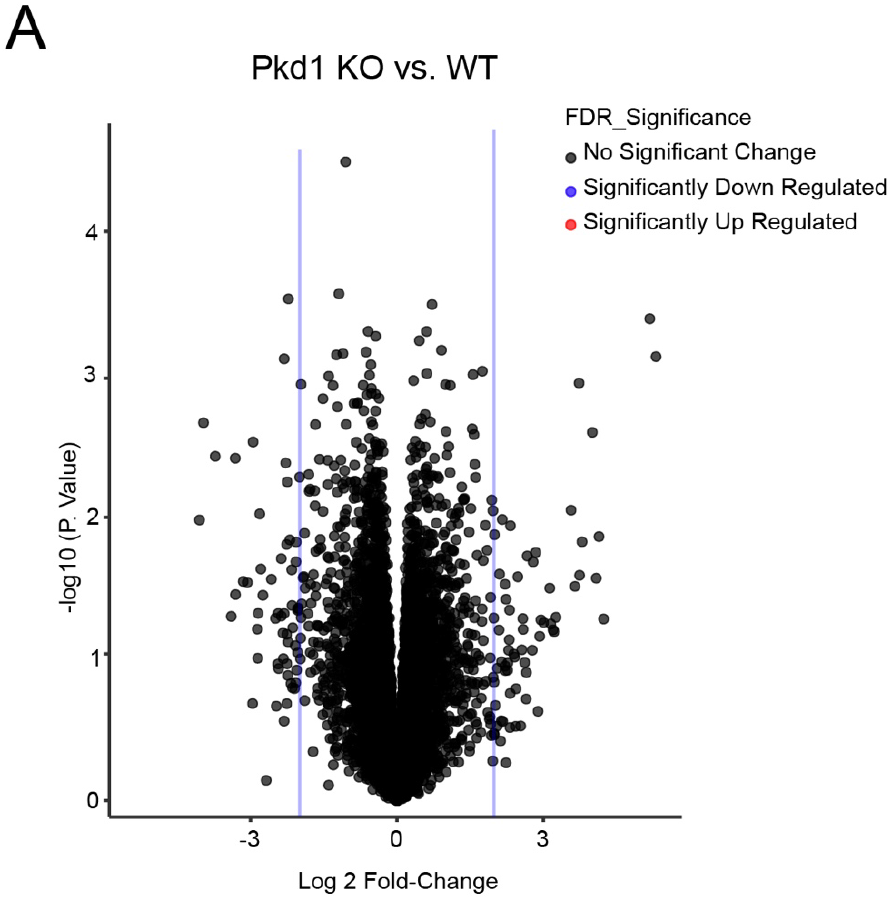
Volcano plot of DEGs of Hoxb7-Pkd1-KO kidneys and WT at E13.5. (A) Volcano plot of differentially expressed genes between Hoxb7-Pkd1 KO kidneys and WT at E13.5. No significant DEGs passed both the >2 log fold-change and p-value < 0.05 thresholds in Pkd1-KO embryonic kidneys compared with WT. N = 3 mice per sample.

**Supplementary Figure 4.**
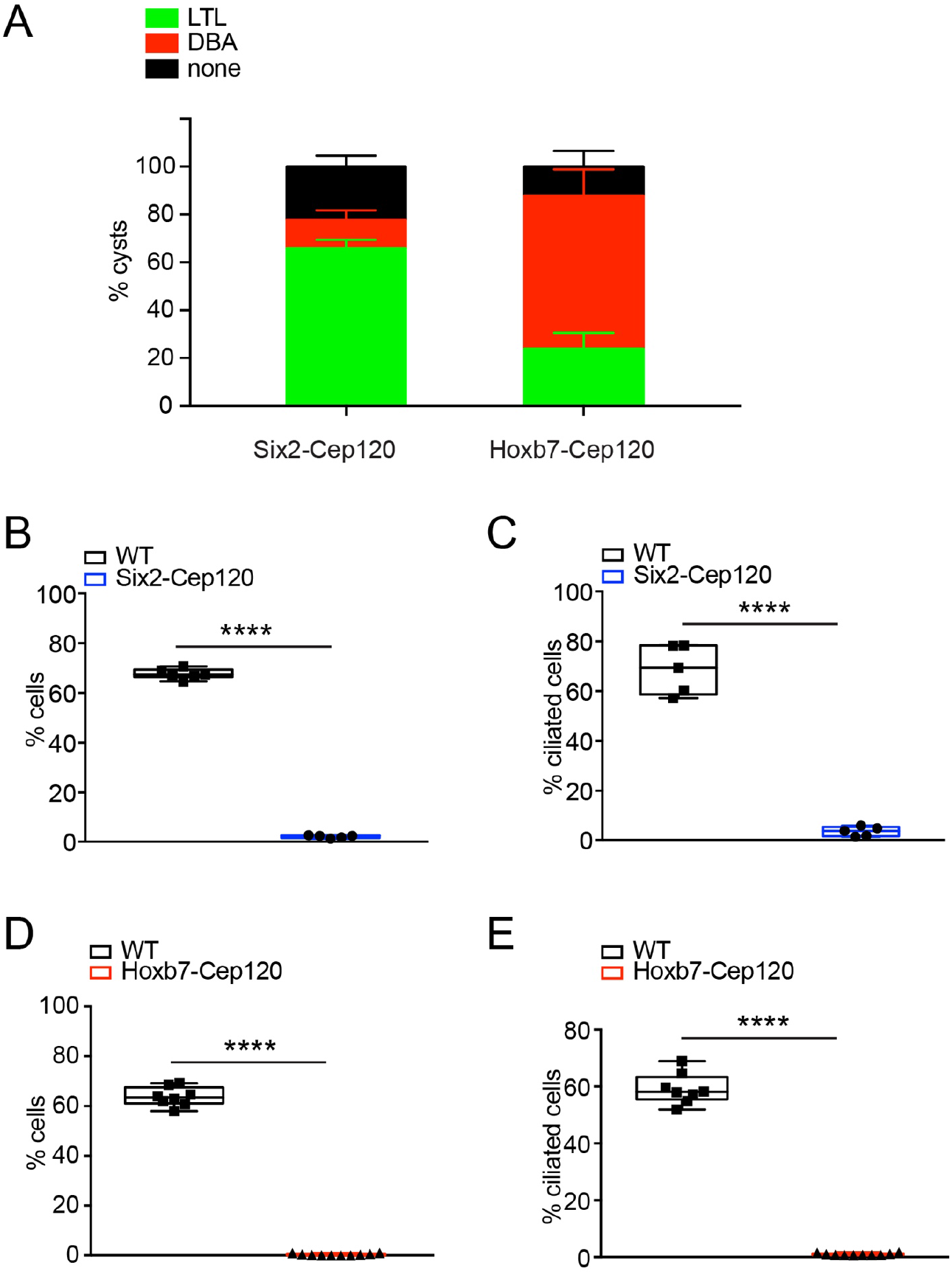
Quantification of centrosome-cilia abundance and cyst origin in postnatal Cep120 knockout mice. (A) Quantification of the percentage of cysts stained for markers of proximal tubules (LTL) or collecting ducts (DBA) in kidney sections of Six2-Cep120 and Hoxb7-Cep120 KO mice at P15. (B-C) Quantification of the percentage of cyst-lining cells lacking centrosomes or cilia in Six2-Cep120 KO and control kidneys at P15. N = 6 mice (WT) and N = 5 (Six2-Cep120 KO). (D-E) Quantification of the percentage of cyst-lining cells lacking centrosomes or cilia in Hoxb7-Cep120 KO and control kidneys at P15. N = 8 mice (WT) and N = 10 (Hoxb7-Cep120 KO).

**Supplementary Excel 1: Bulk RNA sequencing data from E13.5 embryonic kidneys**

Tab1: Raw expression data of all genes from Hoxb7-Cep120 KO and WT kidneys.

Tab2: Differentially expressed genes (|log2 fold-change| ≥ 2) of Hoxb7-Cep120 KO vs. WT kidneys.

**Supplementary Excel 2: Bulk RNA sequencing of postnatal cystic kidneys at P15**

Tab1: Raw expression data of all genes from Hoxb7-Cep120 KO, Hoxb7-Pkd1 KO and WT kidneys.

Tab2: Differentially expressed genes (|log2 fold-change| ≥ 2) of Hoxb7-Cep120 KO vs. WT kidneys.

Tab3: Differentially expressed genes (|log2 fold-change| ≥ 2) of Hoxb7-Pkd1 KO vs. WT kidneys.

Tab4: Differentially expressed genes (|log2 fold-change| ≥ 2) of Hoxb7-Cep120 KO vs. Hoxb7-Pkd1 KO kidneys

